# Organismal robustness and resilience influence the ability to cope with cold snaps

**DOI:** 10.64898/2025.12.22.696020

**Authors:** Conor C. Taff, Daniel R. Ardia, David Chang van Oordt, Jennifer L. Houtz, Nicole A. Mejia, Thomas A. Ryan, J. Ryan Shipley, Jennifer J. Uehling, David W. Winkler, Cedric Zimmer, Maren N. Vitousek

## Abstract

Coping with challenges often requires maintaining performance during poor conditions (robustness) and reestablishing normal performance following disturbance (resilience). Trade-offs between performance under normal and challenging conditions may generate variation in stress coping capacity, but there is little empirical data from wild populations. We studied robustness and resilience to temperature in tree swallows (*Tachycineta bicolor*) using a 39-year dataset that included >75,000 hours of behavioral data. Cold snaps had profound and sustained impacts on fitness; effects on nestlings were mediated by the behavioral robustness and resilience of parents. Parental robustness was predicted by developmental cold exposure and current endocrine state. As expected, trade-offs between peak performance and behavioral robustness influenced susceptibility. Selection on trade-offs may generate population differences in vulnerability to changing temperature regimes.

---

Wild animals are routinely exposed to challenging and unpredictable environmental disturbances (*1*). As a consequence, physiological and behavioral stress response systems are fundamental characteristics of life on earth (*2–5*). Yet, despite the critical importance of coping ability, studies regularly find enormous between-individual variation in the ability to cope with challenges (*6–8*). Why does the same environmental challenge, such as a cold snap or food shortage, result in mortality for some individuals whereas others appear unaffected?

Individual differences in resilience and robustness provide mechanistic insights into why some organisms effectively cope with environmental challenges whereas others do not (*9–12*). Resilience describes the return to a prior state after disturbance, while robustness is the maintenance of performance during an ongoing disturbance. These ideas have a long history in biology (*13*), but there has been renewed interest recently in their potential to unify the understanding of coping with disturbance by connecting physiological, organismal, and population scales (*14–16*). Critically, this framework explains individual differences in coping ability as a result of physiological mechanisms shaped by evolutionary trade-offs, and the maintenance of such variation within populations due to selection pressures acting differently across contexts.

Empirically demonstrating the importance of organismal resilience and robustness in the face of environmental change has been difficult (*17*) because it requires: i) establishing clear fitness impacts of ecological stressors in natural populations, ii) characterizing individual coping phenotypes through extensive behavioral and physiological monitoring under both challenging and benign conditions, and iii) detailed tracking of fitness outcomes. Because animals are exposed to increasingly variable and extreme conditions as a consequence of climate change (*18*), individual differences in coping ability that facilitate the evolution of increased robustness or resilience to disturbance may be a key predictor of population persistence. We studied the coping ability of wild tree swallows (*Tachycineta bicolor*) during the breeding season over 39 years, including 11 years of intensive behavioral monitoring during incubation and provisioning using automated nest-box systems.

Tree swallows have evolved a brief, energetically intensive breeding season that reflects an evolutionary trade-off: by synchronizing reproduction with peak insect availability, they maximize reproductive success, but at the cost of increased sensitivity to environmental variability early in the season (*19*). Even under benign conditions, this compressed window demands high parental investment. For example, females lose approximately 13% of their body mass over just six days after chicks hatch (Fig. 1); this pattern likely facilitates more efficient flight during provisioning (*20*, *21*), but it may also increase vulnerability to deteriorating conditions. In contrast, nestlings grow rapidly and reach or exceed adult average mass within 11 days of hatching (Fig. 1B). Nestlings also develop the ability to thermoregulate over the first 2 weeks of life (*22*). Cold snaps that occur during this period are doubly challenging because they simultaneously increase thermoregulatory costs and reduce food availability (*23*, *24*), often resulting in complete nest failure (*19*, *25*).

**Fig. 1.**
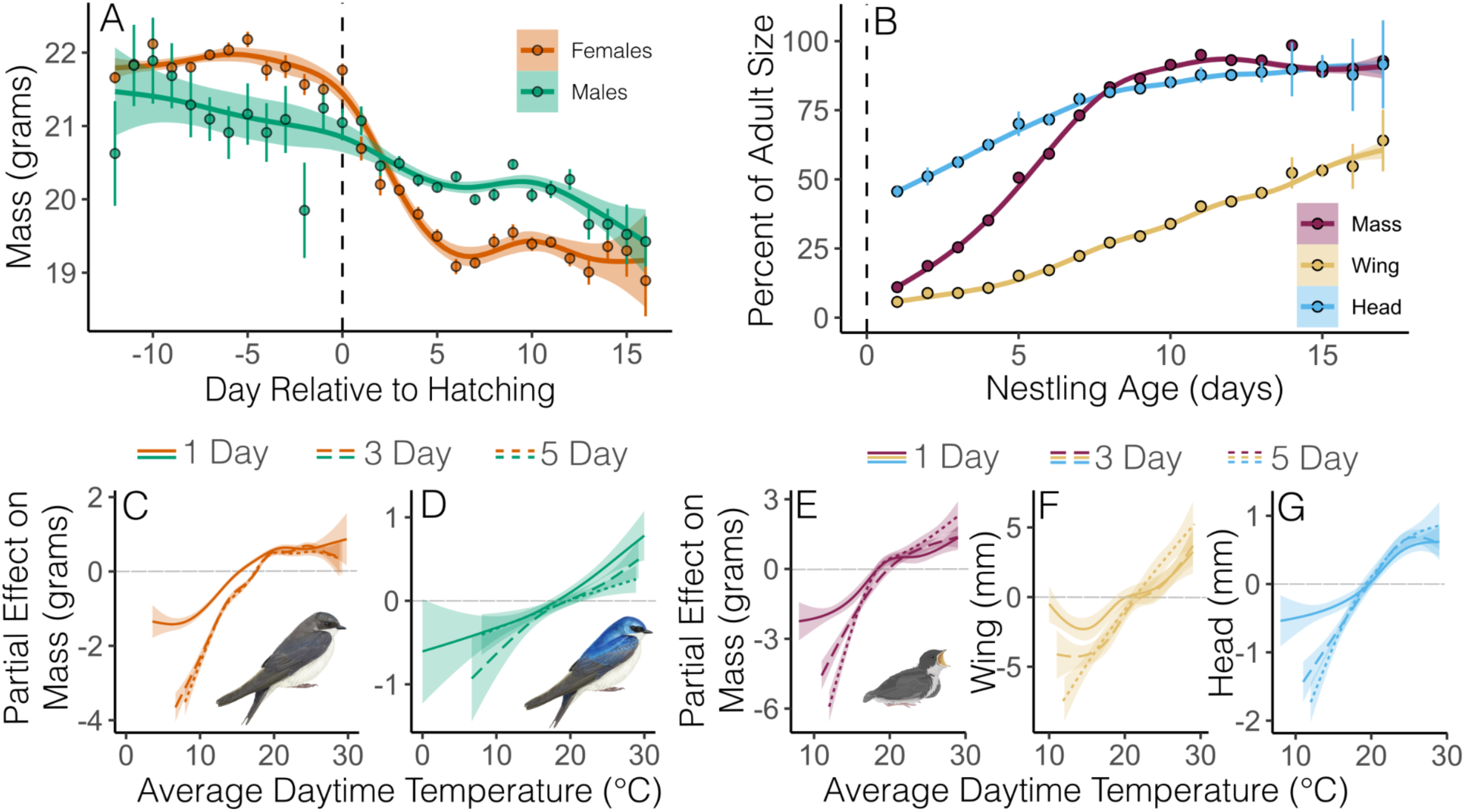
Effects of breeding progression, growth, and temperature on tree swallow mass and morphology. (**A**) Relationship between the day in the nesting cycle and mass of adult females (*n* = 7,843 measures from 3,669 individuals) and adult males (*n* = 2,731 measures from 1,898 individuals). (**B**) Nestling growth as the percent of average adult size attained in mass (*n* = 21,049 measures from 15,983 individuals and 3,506 nests), flattened wing length (*n* = 18,469 measures from 15,099 individuals and 3,315 nests), and head plus bill length (*n* = 15,336 measures from 12,031 individuals and 2,650 nests). In **A** and **B**, points and bars show averages and SEM from raw data, while curves and confidence intervals are plotted from a GAMM. Lower panels show the partial effect of average daytime temperature on (**C**) adult female mass, (**D**) adult male mass, (**E**) nestling mass, (**F**) nestling flattened wing length, and (**G**) nestling head plus bill length from GAMMs that account for day relative to hatching and include a random effect for individual (adults) or nest (nestlings) identity. Adult panels are plotted at the average value for capture day relative to hatching and nestling panels are plotted with age held at 8 days old, during the linear stage of growth. In each panel, average daytime temperature (6 a.m. to 8 p.m.) is shown for the day of capture (solid line; 1 day), the day of capture plus two prior days (long dash; 3 day), and the day of capture plus four prior days (short dash; 5 day). Illustrations by Charlotte Holden, reproduced with permission from the Cornell Lab of Ornithology.

## Breeding season cold snaps reduce mass, growth rate, and survival

After accounting for the breeding stage, cold temperature resulted in lower mass for adult females and males (Fig. 1C-D; Table S1; GAMM smooth term for temperature controlling for day of breeding: females *n* = 7,843, P < 0.001; males *n* = 2,731, P < 0.001). Cold-driven mass loss was most pronounced in females and became more severe with multi-day cold snaps, a pattern not observed in males (Fig. 1C; ΔAIC for female 3-day vs. 1-day model = 713.1).

Nestlings showed similar reductions in mass gain during cold conditions along with delayed structural growth across all ages (Fig. 1E–G; Table S1; GAMM mass *n* = 21,049, P < 0.001; wing *n* = 18,469, P < 0.001; head *n* = 15,336, P < 0.001). Like females, these delays for nestlings were more pronounced when cold conditions persisted for more than one day (Fig. 1E–G; ΔAIC for 3-day vs. 1-day mass model = 303.4; 5-day vs. 1-day wing model = 649.5; 5-day vs. 1-day head model = 385.8). The partial effect of temperature indicated that when temperatures were below 18–20 °C, nestling mass and morphology dropped below the overall average size, closely matching a temperature threshold previously identified as an inflection point for declines in flying insect abundance (*23*).

These direct effects of cold on morphological measurements likely underestimate the full impact of cold snaps, as they do not account for differential mortality or long-term carryover effects of cold-induced growth delays (*26*). At the population level, multi-day cold snaps often cause widespread nestling mortality before measurements can be taken, meaning our assessment excludes many nests that failed entirely due to cold exposure (*19*). At the individual level, we found that the likelihood of successfully fledging increased as average daytime temperature increased when nestlings were 5–12 days old (Fig. S1; binomial model odds ratio of standardized temperature on fledge success = 1.76, 95% CI = 1.58 to 1.95, P < 0.001, *n* = 9,471 nestlings from 1,837 nests). Moreover, we previously showed that nestlings that fledge at a larger mass are more likely to survive and recruit as adult breeders and continue to be relatively large as adults (*27*); thus, the consequences of cold exposure continue into adulthood even for nestlings that do fledge successfully.

Adult females, but not males, that experienced colder conditions during breeding were less likely to return to the site the next year (Fig. S2; female odds ratio of return = 1.10, CI = 1.04 to 1.18, P = 0.003, *n* = 3,673; male odds ratio = 0.94, CI = 0.85 to 1.03, P = 0.17, *n* = 1,863). These results are consistent with the idea that differences in parental investment and body condition during provisioning make females more vulnerable than males during environmental challenges. Together, the immediate and lasting effects of cold snaps during the nesting period highlight the severe ecological challenge posed by low temperatures. Variation in the ability to cope with this ecological challenge may be an important predictor of fitness, especially since nestling exposure to cold snaps has increased as a consequence of earlier breeding due to climate change (*19*).

## Individuals vary in their behavioral resilience and robustness to cold

Given the challenge that cold snaps impose, the ability to cope with cold through behavioral adjustments could generate differences in breeding success. For example, some adults might be able to buffer the costs of cold exposure on nestlings by increased brooding or feeding effort (*28*). We used automated systems to continuously measure behavioral variation in incubation (for females) and provisioning (for both sexes). These behaviors are impacted by environmental temperature and are essential to nestling development and survival (*29*).

In addition to monitoring variation in the response to temperature, we also used these systems to record recovery time after the disturbance imposed by capture. This measure of resilience is distinct from sensitivity to cold and allowed us to assess whether individual variation in resilience generalized across contexts (*14*). In total, we extracted a suite of 16 behavioral traits for each nest that we characterized as indicative of mean individual differences (e.g., incubation on-bout length), robustness to challenging conditions (e.g., slope of feeding rate to declining temperature), or resilience to disturbance (e.g., differences in the recovery to normal behavior one day after experiencing low temperature; full description of each behavior in Table S2).

At the population level, the length of incubation on- and off-bouts increased at both cold and hot ambient temperatures with an inflection point around 19°C (Fig. 2A). Incubation-bout characteristics had a profound effect on the thermal environment experienced by developing embryos. Longer on-bouts resulted in a higher maximum egg temperature and a greater temperature increase during the bout (Fig. S3; Table S3; LMM with *n* = 44,220 bouts from 209 nests, bout duration on max temperature standardized β = 0.08, 95% CI = 0.07 to 0.08, P < 0.0001; duration on temperature swing β = 0.21, CI = 0.21 to 0.22, P < 0.0001). Similarly, longer off-bouts resulted in lower minimum egg temperature and a greater decrease in temperature during the bout (Fig. S3; Table S3; LMM with *n* = 44,834 bouts from 209 nests, bout duration on minimum temperature standardized β = -0.09, CI = -0.09 to -0.08; duration on temperature swing β = -0.20, CI = -0.21 to - 0.19, P < 0.0001). Both of these patterns were especially pronounced when ambient temperature was low (Fig. S3; Table S3; interaction between ambient temperature and bout duration P < 0.0001). Feeding rate was also strongly influenced by ambient temperature, with the highest hourly feeding rate for females and males at 25°C and declining feeding rate at lower and higher temperatures, but with the lowest feeding rates observed during cold conditions (Fig. 2B). Reduced feeding rate directly impacts the amount of nutrients that nestlings receive during the critical growth period (*30*).

**Fig. 2.**
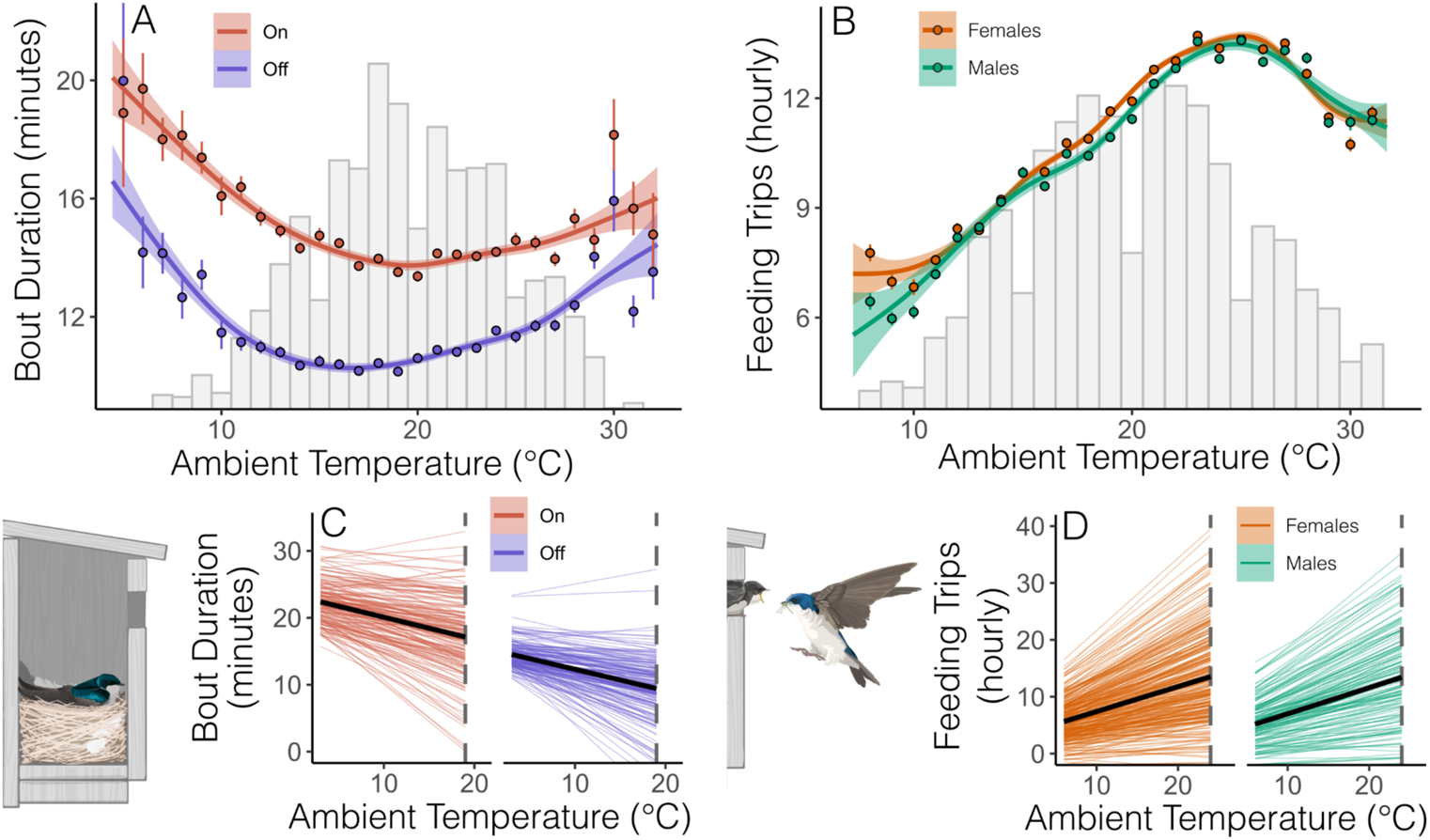
Effect of temperature on behavior at the population level and between individual variation in sensitivity to declining temperature. (**A**) Variation in the duration of incubation on bouts and off bouts by temperature (*n* = 44,361 on bouts and 44,987 off bouts over 17,258 daytime hours from 209 nests and 184 unique females). (**B**) Variation in hourly daytime provisioning rate for adult females and males by temperature (*n* = 1,020,830 feeding trips over 61,142 hours from 376 nests attended by 314 unique females and 206 unique males). In **A** and **B** lines and confidence intervals are from a GAMM and points show the mean and SEM of raw data for each temperature value; histograms illustrate the distribution of observed hourly ambient temperature in the dataset. (**C** & **D**) Random effect estimates for each nest illustrating variation in the linear response to declining temperature below 19 ℃ for incubation (**C**) and below 24 ℃ for provisioning (**D**). Solid black lines show the population average response. Illustrations by Charlotte Holden, reproduced with permission from the Cornell Lab of Ornithology.

At the individual level, we found variation reflected in overall bout length and feeding rate (between-individual differences in mean behavior), in the sensitivity of those behaviors to declining temperatures (differences in robustness), and in their recovery from cold temperatures experienced the previous day (differences in resilience; Fig. 2 C-D; Table S4). Overall, between-individual differences in mean behavioral level explained more variation than differences in resilience or robustness to temperature (Table S4; random intercept explained 7-24% of variation and random slope explained 0.3-4.2% of variation), but including random slopes for individuals improved the fit of each model (Table S4; random intercept vs. random slope models ΔAIC range = 18.8 to 1,040.4). Separately from the impacts of cold exposure, we also found substantial variation between individuals in resilience to general disturbance reflected in the amount of time required to return to the nest after an experimental capture and the degree to which afternoon feeding rate was impacted by morning captures (Fig. S4).

Across the 16 behavioral traits we examined, we found extensive correlations indicative of both integrated behavioral phenotypes and trade-offs among components of the behavioral response to cold. Several clusters of correlated traits emerged (Fig. 3; Fig. S5). First, we observed variation in incubation strategies: females with longer on-bouts also tended to have longer off-bouts, suggesting consistent individual differences in incubation style. Second, provisioning rates and temperature sensitivity were closely aligned within and between sexes. This pattern may reflect coordinated investment in parental care between mates and/or shared aspects of individual quality (*31*) and suggests that coping ability and behavioral investment are shaped not only by individual traits but also by the social mate.

**Fig. 3.**
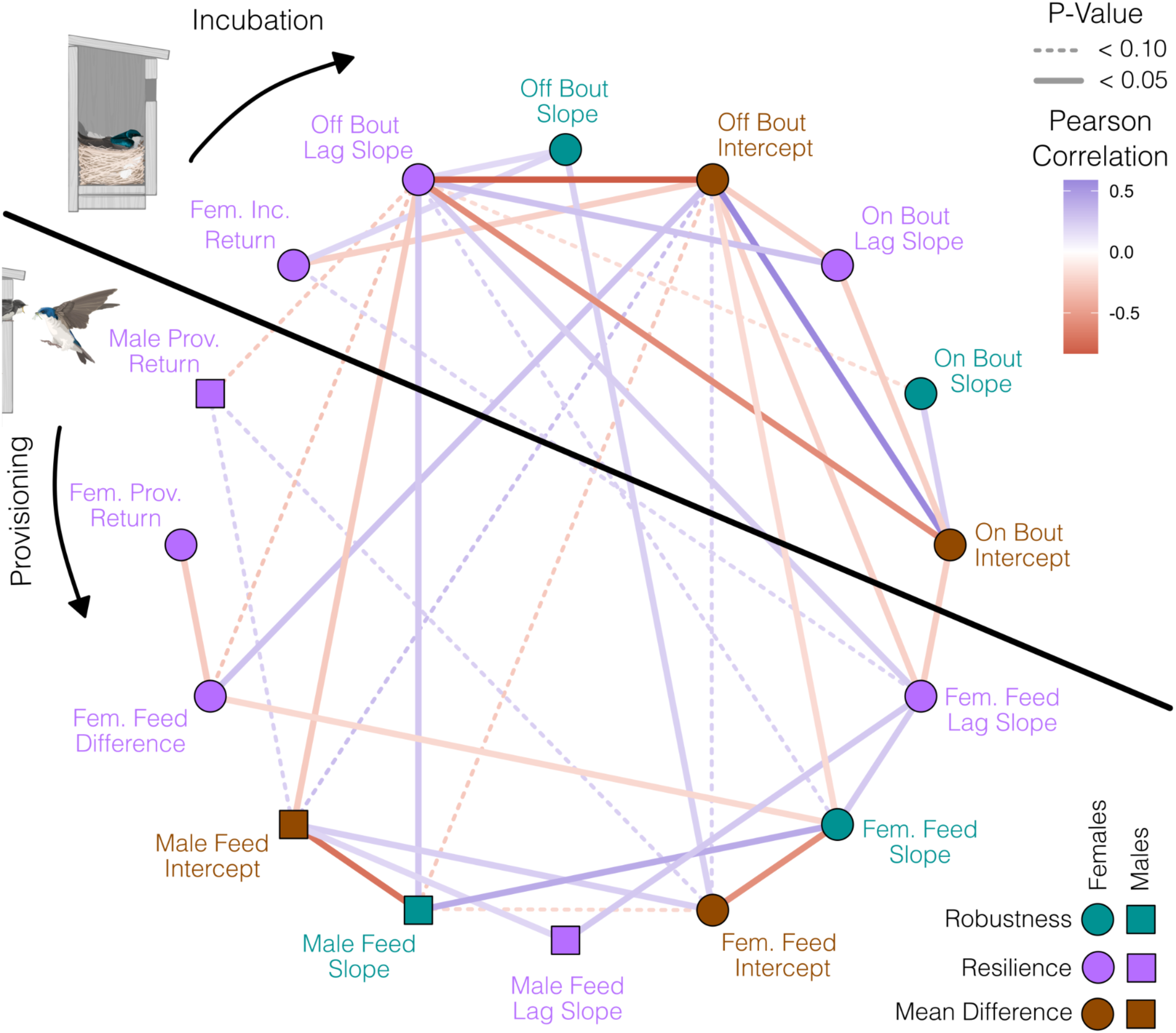
Correlation between behavioral characteristics extracted from automated behavioral monitoring systems. Pairwise correlations that were significant (solid lines; P < 0.05) or non-significant trends (dashed lines; P < 0.10) are included. Colors indicate whether each behavior is related to robustness, resilience, or mean individual differences and shapes differentiate males from females. The color gradient of connection lines indicates the correlation coefficient of each pairwise correlation. When necessary, measures are inverted so that higher values always indicate increasing resilience, robustness, and mean behavioral output. Full descriptions of each behavior are given in Table S2. Illustrations by Charlotte Holden, reproduced with permission from the Cornell Lab of Ornithology.

We also found that coping styles were reflected as correlated sets of robust or resilient behaviors across different contexts (*32*). In general, distinct measures of resilience tended to be positively correlated, whereas between-individual differences in average incubation or provisioning levels tended to be negatively correlated with resilience (Fig. 3). This pattern is predicted by several models of coping ability, which suggest that one cost of high levels of peak performance, such as high average feeding rates or long on-bouts, might be increased fragility that leads to lower robustness or resilience of those same traits when challenging conditions occur (*9*, *17*, *33*).

## Behavioral differences are associated with fitness outcomes

Parental investment in both incubation and feeding had direct impacts on nestling outcomes. Longer incubation on-bouts were associated with a higher chance of hatching (Fig. 4; Table S5; odds ratio = 1.92, 95% CI = 1.01 to 3.27). The effect of incubation behavior also extended into the nestling period, as longer on-bouts were associated with higher survival from hatching to day 12 (odds ratio = 1.85, CI = 1.07 to 2.52). This finding might reflect carryover effects from incubation or continued differences in female parental investment, such as increased brooding behavior. Increasing on-bout length from 1 sd below to 1 sd above the mean was associated with a 19% and 48% increase in hatching and survival to day 12, respectively. The effects of incubation waned as nestlings aged and there was no relationship with later survival, recruitment, or morphological measurements and no evidence that sensitivity of on-bout length to temperature was related to nestling survival or growth (Fig. 4; Table S5-6).

**Fig. 4.**
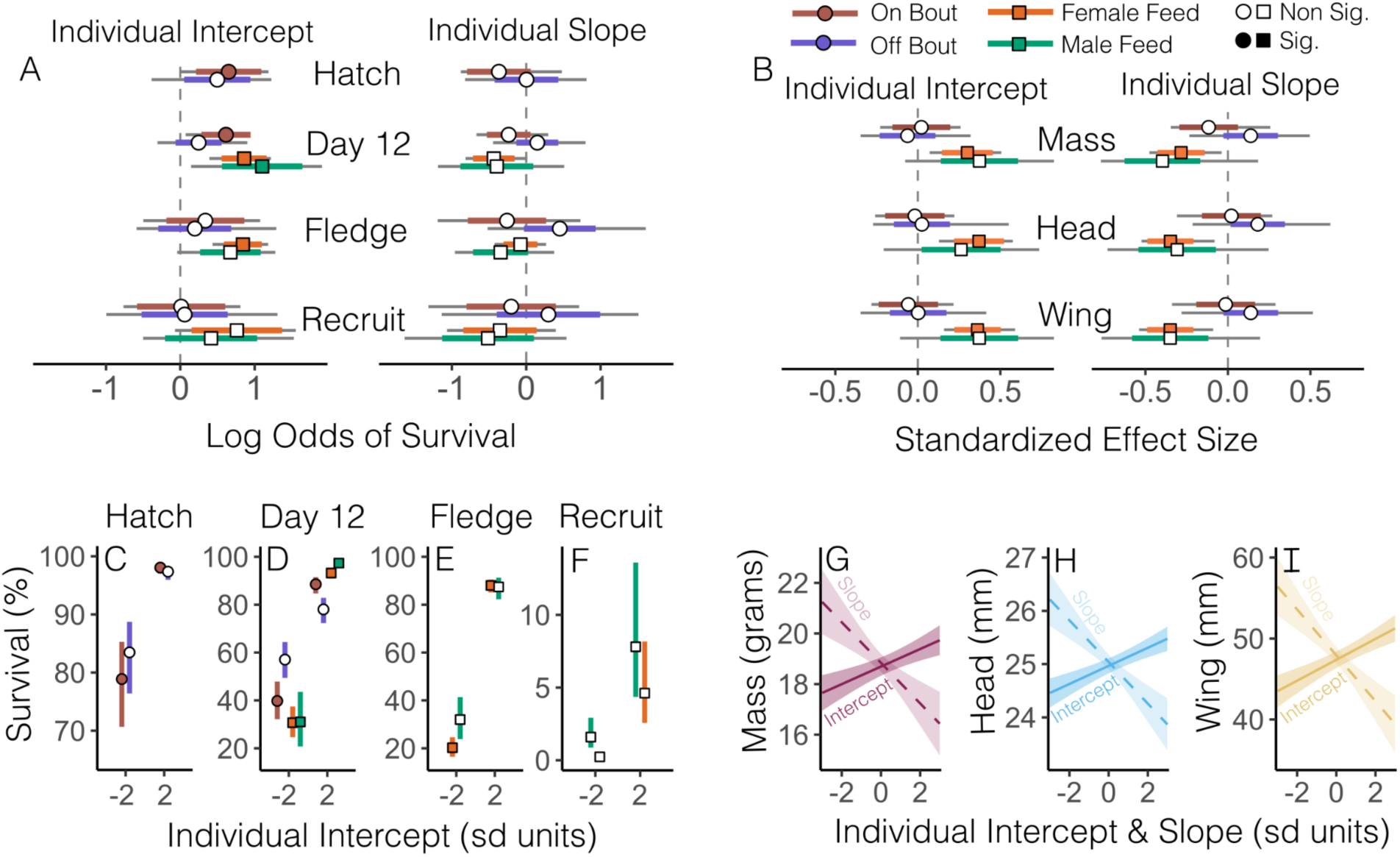
Relationships between adult incubation and provisioning behavior and nestling survival and growth. (**A**) Estimated log-odds from binomial models of surviving from laying to hatching, from hatching to day 12 or to fledging, and from fledging to recruiting in a subsequent year with the random intercept and slope of on bout length, off bout length, female feeding rate, or male feeding rate as predictors. (**B**) Standardized effect sizes for models with average nestling mass, head, or flat wing on day 12 with the intercept and slope of behavioral measures as predictors. Individual intercepts and slopes are best linear unbiased predictors (BLUPs) for random effects of individual identity in each model. In A and B, colored bars show 95% confidence intervals from models and gray bars show confidence intervals from 100 bootstrap iterations to account for uncertainty in BLUP estimation. Filled shapes indicate effects in which the bootstrap confidence interval did not cross zero. (**C** to **F**) Point estimates and confidence intervals for predicted survival percentage from nests with intercept BLUP estimates 2 sd below and above the mean for (**C**) hatching, (**D**) day 12, (**E**) fledging, and (**F**) recruiting. (**G** to **I**) Model predicted relationships between the BLUP for female feeding intercept (solid line) and slope (dashed line) compared to average nestling (**G**) mass, (**H**) head, and (**I**) wing. Sample size varies by model; see Table S6-7 for full details.

Variation in feeding behavior had more pervasive effects on both survival and nestling growth. Female feeding rate was positively associated with survival to day 12 (Table S5; odds ratio = 2.36, CI = 1.48 to 3.38) and survival to fledging (odds ratio = 2.32, CI = 1.54 to 3.26); nestlings with mothers who fed at high rates were also more likely to recruit as breeding adults, although the confidence interval for this effect crossed one (odds ratio = 2.14, CI = 0.93 to 4.72). Greater sensitivity of feeding rate to declining temperature also tended to be associated with reduced survival to day 12, although the effect was not significant (Table S5; odds ratio = 0.65, CI = 0.44 to 1.01). As with incubation, the effect sizes of feeding behavior on survival rates were large, such that an increase in feeding rate from 1 sd below to 1 sd above the mean was associated with a 68% increase in the likelihood of fledging.

In addition to the effects on survival, higher female feeding rate was correlated with increased average nestling mass, head size, and wing length on day 12 (Fig. 4; Table S6; feeding intercept on mass β = 0.3, CI = 0.07 to 0.51; head β = 0.37, CI = 0.13 to 0.58; wing β = 0.36, CI = 0.16 to 0.59). In contrast, females that reduced their feeding rate more steeply with declining temperature produced nestlings that were smaller in all three measures on day 12 (Fig. 4; Table S6; feeding slope on mass β = -0.28, CI = -0.48 to -0.04; head β = -0.35, CI = -0.52 to -0.08; wing β = - 0.35, CI = -0.54 to -0.09). Because feeding slope and intercept were positively correlated, their opposite effects on nestling growth likely reflect a trade-off in feeding behavior. Females that were able to feed at a high rate, but limit sensitivity to temperature given their feeding rate, produced the largest nestlings with the greatest chance of survival and recruitment. The qualitative patterns and effect size for male feeding rates were similar to females, but with a smaller sample size the bootstrapped confidence interval for male effects crossed zero (Fig. 4).

## Early life conditions and current physiology influence behavioral sensitivity

Understanding the proximate mechanisms that generate differences in behavioral robustness and resilience is critical to predicting how resilience will evolve in the context of climate change (*34*, *35*). We previously found that early life conditions–particularly temperature experienced during the late incubation stage–influence the physiological stress response of adult tree swallows (*36*). In accordance with those results, average temperature during late incubation of the hatch year was related to female feeding rate sensitivity to temperature as adults. Females that were incubated during colder conditions were less sensitive to declining temperatures and as a result provisioned their own nestlings at higher rates when temperatures dropped (Fig. 5A; Table S7; *n* = 54 females, temperature by hatch year incubation temperature interaction F = 17.62, P < 0.0001). In contrast to the detrimental effects of cold temperature for 8 day old chicks described above, this result suggests that exposure to cold temperatures during an earlier critical period of development primed individuals to respond more effectively to later life cold snaps. Crucially, earlier breeding due to climate change at this site has increased the likelihood of exposure to cold temperatures during the vulnerable nestling growth period, but not the incubation period (*19*). Because relatively few nestlings from our site return as adults, we did not have a sufficient sample size to examine early life effects on male behavior or on incubation behavior.

**Fig. 5.**
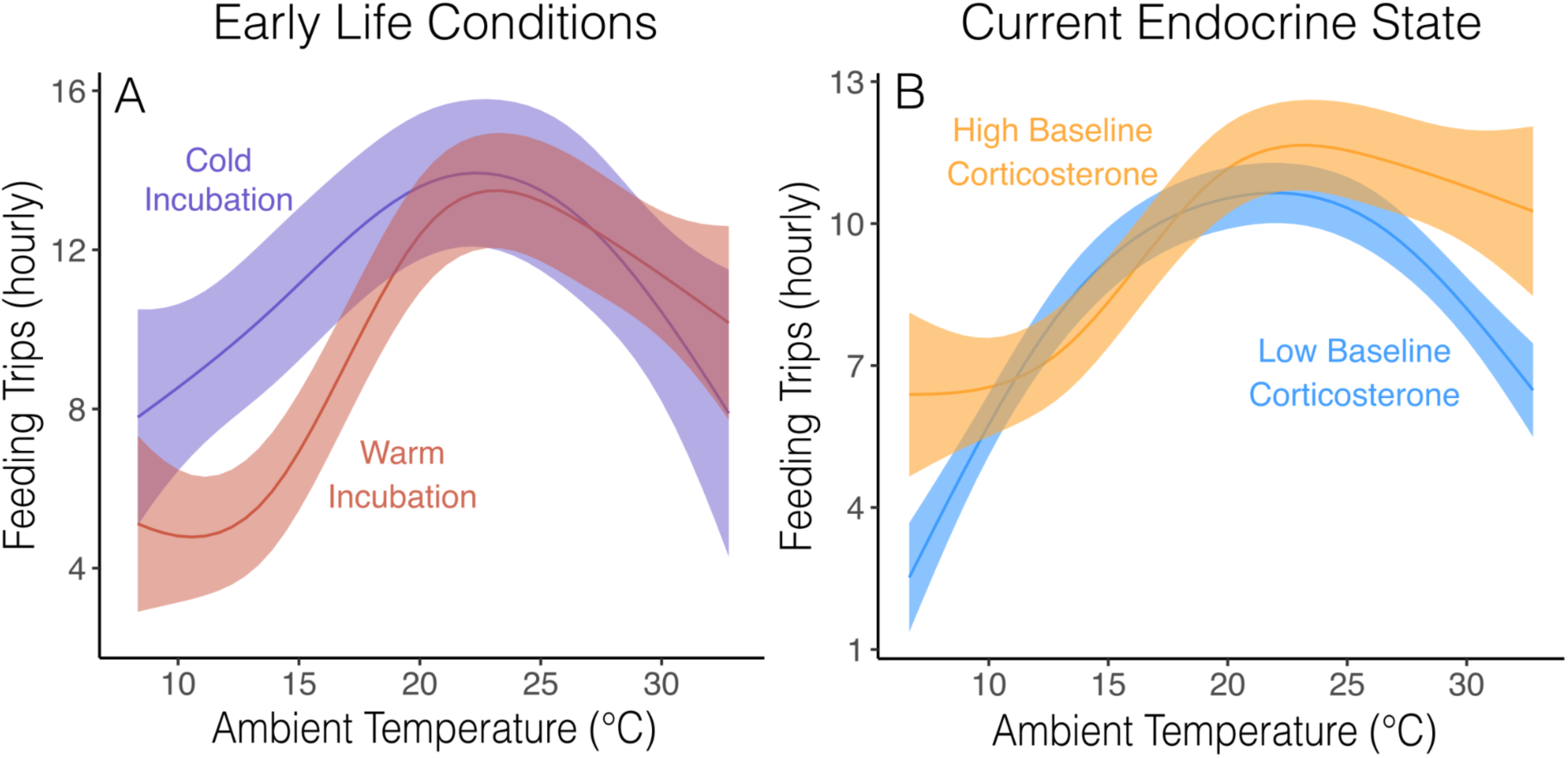
Interaction between current temperature and **(A)** the average temperature during late incubation of the hatch year or **(B)** baseline corticosterone during the early provisioning period. Predictions are extracted from GAMMs with continuous interactions and plotted at the 5th and 95th percentile for early life temperature and baseline corticosterone to illustrate the interaction. Models included a random effect controlling for female identity.

We also found evidence that endocrine state mediates the behavioral response to temperature. Glucocorticoid regulation plays a critical role in both energy allocation through variation in baseline concentrations and in the response to acute challenges through rapid activation of the corticosterone stress response (*3*, *4*). We found no evidence for an association between baseline or stress-induced corticosterone and the incubation response to temperature (Table S7). However, feeding rates for females with higher baseline corticosterone were less sensitive to temperature fluctuations and, as a result, these females fed nestlings at higher rates during extreme temperature conditions (Fig. 5B; Table S7; *n* = 147 females, temperature by baseline corticosterone interaction F = 30.23, P < 0.0001). We found no evidence for an interaction between stress-induced corticosterone and the feeding rate response to temperature fluctuations (Table S7).

Although elevated baseline corticosterone is often associated with reduced fitness (*37*), normal fluctuations in baseline corticosterone play an important role in metabolic regulation and can support breeding during demanding conditions (*4*, *38*) and moderate elevation is therefore often associated with higher reproductive investment and metabolic rate (*39*, *40*). Our results are consistent with the idea that higher maternal investment associated with elevated baseline corticosterone might buffer nestlings from inclement conditions. In previous analyses that did not account for temperature, we found that corticosterone-fitness relationships are context-dependent in this population (*41*). Taken together, our results suggest that early life conditions and physiological state might mediate how effectively individuals can cope with conditions encountered during the breeding season.

## Conclusions

Obligate aerial insectivores, such as tree swallows, have declined precipitously over the past 50 years (*42*), but the declines have been geographically uneven. At our study site, tree swallows have advanced their breeding sufficiently to track average temperature during the egg laying period. However, early season temperature is more variable and, as a consequence, nests are more likely to be exposed to cold snaps during the most vulnerable period of chick growth than they were historically (*19*). Variation in behavioral resilience to cold and the ability of resilience to evolve with increasing exposure to variable temperatures is likely to be an important predictor of population persistence in the face of climate change.

We found strong evidence that multi-day cold snaps have dramatic consequences for tree swallows, that adults differ in their behavioral robustness and resilience to declining temperature, and that these differences predict fitness outcomes. Our results suggest that some individuals possess integrated behavioral phenotypes that make them more capable of coping with an ecologically relevant challenge. However, we also found that early life conditions played a role in shaping adult phenotypes. Thus, prior exposure and experience might be an important determinant of coping ability (*43*) and population vulnerability could depend on the interactive effects of changing climate conditions during both development and adulthood. Our work demonstrates the utility of the resilience and robustness framework for understanding the ability to cope with challenges.

## ACKNOWLEDGMENTS

We thank the many members of the tree swallow research team who contributed to data collection over the years of this project. We receive helpful feedback on a draft of this manuscript from Anurag Agrawal, Mike Webster, and members of the Vitousek Lab.

## AUTHOR CONTRIBUTIONS

All authors contributed to data collection. C.C.T. designed the study, conducted the analyses, and led the writing with input from M.N.V and J.R.S. All authors contributed to the writing of the final manuscript.

## DATA & CODE AVAILABILITY

All data and code required to produce the results presented here have been publicly archived on Zenodo (code DOI: 10.5281/zenodo.15864802; data DOI: 10.5281/zenodo.15864830).

## SUPPLEMENTARY MATERIALS

Materials and Methods

Figs. S1 to S6

Tables S1 to S7

References (*44-52)*

## METHODS

### General field and lab methods

#### Population monitoring and sampling

We monitored tree swallows in nest boxes in Tompkins County, New York from April to July 2013 to 2024. Tree swallows at this location have been studied continuously since 1986 and for some analyses we also drew from the full 39 years of observation. During the last decade, each nest was typically checked every other day throughout the breeding season and every day around the projected hatching date. After banding on day 12, nests were avoided to prevent forced fledging and checked a final time on day 24 to determine fledging success for each banded nestling. This schedule allowed us to determine nestling survival and the dates of key events as well to schedule sample collection and measurements identically across all nests.

Adult females were captured in the nest box on day 6–8 after the onset of incubation (early incubation) and again on day 2–8 after hatching (provisioning). In some years, females were also captured 1–2 days before the projected hatching day (late incubation). All female captures occurred between 6am and 10am. Males were captured opportunistically at a subset of nests 2–8 days after hatching. Nestlings were measured on day 12 after hatching between 12pm–3pm. If a bird had not been previously captured, we applied a unique USGS aluminum band for identification. For adults, we also added a passive integrated transponder (PIT) to collect provisioning data (see below). For each bird, we measured mass, flattened wing length, and the length from the back of the skull to the tip of the bill (referred to as head size).

For most adults, we also collected blood samples (<70 μl each) to assess the corticosterone response to a standardized handling restraint protocol. This included one sample taken within 3 minutes of disturbance to measure baseline corticosterone and a second sample taken 30 minutes after capture to measure stress-induced corticosterone. Plasma was separated and frozen within 3 hours of capture and used to measure corticosterone concentration. Corticosterone was measured following a triple ethyl acetate extraction using an ELISA kit (DetectX 4-H; Arbor Assays) that has previously been validated in this population (*44*).

Nests in this population have been the subject of a wide range of experimental studies. In recent years, those have included hormone, temperature, plumage color, and predator manipulations. For the purposes of this study, we only included nests that were not part of any experiment or that received either a control or sham control treatment in the experiments described above. For analyses using the longer term dataset, we excluded any nests that were subject to major manipulations or selected only data that was collected before any manipulations occurred.

#### Automated behavioral data collection

We focused our behavioral analysis on a subset of nests that had equipment installed to enable continuous behavioral data collection. First, to collect incubation data, we used a temperature logger (HOBO data logger UX100-014M; Onset) with a wired thermocouple embedded in a fake tree swallow egg. The fake egg was anchored in the nest cup amid the real eggs and logged the temperature every 10–30 seconds. This setup allowed us to determine the start and end of on-bouts and off-bouts as indicated by the change in temperature of the fake egg. We used a custom script along with manual verification to process the raw data stream for each nest into behavioral incubation data (see below: thermocouple data processing).

Second, to characterize provisioning behavior, we installed a radio frequency identification (RFID) device at each nest box with an antenna loop around the nest box entrance hole (*45*). This unit was programmed to record the identity of PIT tags at the nest box continuously from 6am to 8pm every day with a timestamp for each record. We used a previously described custom script to convert this data stream into the number of hourly provisioning trips made by the female and (when available) the male at each nest (described in *46*). Briefly, this script identified unique feeding visits from the raw record of RFID identifications based on the time between reads and was calibrated with video observations of feeding behavior across the nestling growth period. Running raw data through the script resulted in an output with one row per hour of observation along with a female and male feeding rate for that hour.

#### Thermocouple data processing

The complete script to process thermocouple files is included with detailed annotation in the archive that goes with this manuscript. At a high level, the goal of the processing script was to take a stream of temperature values recorded every 10–30 seconds and identify break points representing the beginning of on and off bouts. We accomplished this by first reading in each file and generating a locally estimated scatterplot smoothed trace of the temperature record tuned to capture the rise and fall of bouts while filtering out abrupt jumps in temperature. Next, we developed an algorithm that used the smoothed temperature trace to identify the proposed start of declining periods (off-bouts) and increasing periods (on-bouts). For this task, we coded several tuning parameters that determined the flexibility of the fit line, the minimum temperature change required in a bout, and the minimum temperature level to count as an on bout. We adjusted these values manually to achieve a fit that worked well for our population. This algorithm produced an output with one row per bout indicating an on vs off bout, the duration, the starting temperature, and the ending temperature.

Next, we ran each thermocouple file through the algorithm described above in a batch and produced a series of trace plots for each nest (examples shown in Fig. S6). Trace plots showed the raw data, the smoothed trace, and the proposed start and end points of each bout. While the algorithm worked very well for clear bouts, we found that there were many sections that did not record clear bouts. We could not always identify why a reading was bad, but these likely resulted from a variety of reasons such as an unplugged or malfunctioning thermocouple, low battery, or poor thermocouple placement within the nest. In order to be confident that only good data were included in our behavioral scores, we manually scanned every trace file and compiled a list of sections to be excluded based on start and end times from the trace files.

Finally, we merged the exclusion file with the data output and excluded any bouts that had been scored during an excluded section to arrive at a final data set. For downstream processing, we also filtered out any information from nighttime records (females typically incubate all night). While the final dataset is a clean record of on- and off-bout timing, we are generally less confident in the absolute temperature of the egg thermocouple because small differences in placement might impact absolute temperature (e.g., between vs. on the outside edge of the real eggs). Thus, we present some analyses of absolute temperatures at the aggregate level, but for analyses of individual differences we focus on timing of on- and off-bouts.

### Data Analysis

#### Seasonal changes in mass and growth and the effect of temperature on mass and survival

We merged long term banding data (1986–2024) with ambient temperature data to assess the impact of temperature on body mass (adults and nestlings), growth (nestlings), and survival (adults and nestlings). Hourly temperature records were obtained from the Ithaca Airport Weather Station (1996–2024) and from the Game Farm Road Weather Station (prior to 1996) from the Northeast Regional Climate Center at Cornell. Both weather stations are within 5km of the observed nests.

Over the course of the 38 years of observation, adults have been captured and measured every day of the breeding cycle and across a range of nestling ages. This variation in sampling timing was crucial, because it allowed us to characterize the expected change in mass across the season for adults and the expected growth pattern with age in nestlings. Female tree swallows undergo a seasonal decline in mass, which is likely adaptive (*20*, *21*), as they progress from incubation into provisioning. These patterns are important when considering temperature effects because we need to account for the non-linear change in morphology and mass separately from any changes associated with temperature.

After plotting overall seasonal patterns for descriptive purposes, we used generalized additive mixed models (GAMMs) to ask how temperature impacts population level morphological measurements. We used GAMMs rather than a linear approach because we expected responses to temperature to be non-linear and threshold dependent, as has previously been shown for flying insect abundance at this location (*23*). For all GAMMs, we restricted the number of basis dimensions (k) to a maximum of 5 in order to maintain biological meaning and prevent overfitting. We expected temperature to have a stronger effect on mass and growth when cold conditions persisted for a longer time. Therefore, we fit models with average daytime temperature (6 am to 8 pm) on three timescales: i) the morning of capture; ii) the morning of capture plus the two prior days; iii) the morning of capture plus the four prior days.

In each model, we included mass, wing length, or head size as the response variable. Predictors included a smoothed fit for day relative to hatching and a smoothed fit for one of the three temperature windows. We plotted the predicted effect of temperature while holding adult panels at the average capture day and holding nestling panels at 8 days old, during the linear growth phase. We assessed the support for models using different temperature time scales by comparison of AIC values.

To assess the impact of temperature on survival, we fit generalized linear mixed models with a binomial distribution. We focused on average temperature from nestling ages 5–12 days old because this period has been identified as a particularly vulnerable time for cold snaps to occur because nestlings are in the process of developing the ability to thermoregulate (*19*). While adults have less difficulty thermoregulating, cold temperatures at this age are challenging because it is difficult to forage for enough food and provide enough warmth from brooding to keep nestlings alive. In each model, we included temperature as a standardized predictor and nest or individual identity as a random effect as appropriate.

#### Variation in behavioral sensitivity to challenges

We followed a similar approach to modeling individual differences in behavioral sensitivity for incubation data (per bout duration) and provisioning data (per hour feeding rate). While individuals might differ in the full functional shape of the behavioral response to a range of temperatures, we were most interested in the response to cold temperature and fitting linear mixed models with individual random effects is a much more tractable approach than estimating individual level non-linear curves. Therefore, we restricted the observations for these analyses to temperatures below the point at which the aggregate population response began to increase (bout duration, 19 °C) or decrease (feeding rate, 24 °C). In this range, the population response was approximately linear and we were able to use linear mixed models with random intercepts and slopes for each individual.

Although we realize that other factors likely contribute to behavioral variation, the goal of these analyzes was to focus exclusively on the main response to declining temperature and we therefore kept the structure as simple as possible. We initially fit four models with either on-bout duration, off-bout duration, female hourly feeding, or male hourly feeding as the response variable. Predictor variables included hourly ambient temperature coinciding with the observation period and a random intercept and slope for individual identity. In cases where the same individual was observed in multiple years, we randomly excluded all but one year of observation. These models yielded measures of overall incubation or feeding level from random intercepts (mean differences) and individual measures of sensitivity to declining temperature from random slopes (robustness). We extracted the best linear unbiased predictors (BLUPs) for random effects and used them in downstream analyses (see below).

We next fit a similar set of models focused on recovery from prior cold exposure (resilience). These models included the same four response variables and current temperature, but added a predictor for average temperature on the day prior to observation along with an individual level random slope for the response to the prior day’s temperature. From these models we extracted the individual slope to average temperature the day before observation as a measure of how impacted individuals were by recent temperature while controlling for current temperature (resilience).

In addition to these model based behavioral measures, we also used RFID records to extract two additional measures of resilience that were not directly related to cold exposure as a way to assess the consistency of resilience across contexts. First, we recorded exact release times at every capture and used RFID records at the nest box to record the latency to return to the box and resume normal breeding behavior. This measure was collected during both incubation and provisioning for females (because they were typically captured twice per season) and during provisioning for males. Second, for females we also measured the degree to which a major disturbance (early morning capture) impacted feeding rate as a measure of behavioral resilience to disturbance. For this calculation we compared the average hourly feeding rate from 1–4pm on the day before capture to the afternoon of capture. Females that had reduced feeding rates were interpreted as being less resilient.

In total, this approach yielded 16 distinct behavioral measures. We used simple pairwise correlations to describe relationships between these behaviors. Sample sizes varied for each measure and we included all available data for each pair-wise comparison. When necessary, we inverted behavioral measures such that larger values always indicated increased output, increased robustness, or increased resilience. The goal of this approach was a qualitative description of links between different measures of mean behavioral output, robustness, and resilience, rather than a strong test of a narrow hypothesis and we therefore illustrated both significant correlations and non-significant trends along with correlation coefficients.

#### Early life conditions

Most tree swallows born at our site disperse to other nearby locations if they survive to breed as adults (*47*). Moreover, many returning adults in the last decade were entered into manipulative experiments or did not have the monitoring equipment installed to collect behavioral data. Despite these limitations, we identified 54 females that were born at our site, returned as adults, were not manipulated, and had provisioning data available. Unfortunately, few of these females had incubation data available and we focused only on provisioning effects. For these females, we calculated the average daytime temperature during the late stages of their natal incubation (1 to 5 days before hatching). We focused on this time period because it was previously identified as an important period for influencing long term endocrine phenotype (*48*).

We then fit a generalized additive mixed model (GAMM) using the ‘gamm’ package in R (*49*) with a tensor product smooth interaction between current temperature and natal incubation temperature on feeding rate with female identity as a random effect. We assessed support for this model by comparing the full model to reduced models that included individual smooth terms for current and natal temperature or only current temperature by AIC values. In order to facilitate interpretation of the continuous interaction, we plotted the predicted relationship between current temperature and feeding rate for females in the 5^th^ and 95^th^ percentile of natal temperature exposure.

#### Current endocrine state

We used GAMMs to ask whether baseline or stress induced corticosterone measured during incubation and provisioning was related to incubation bout length and feeding rate, respectively. For each response (on-bout length, off-bout length, and female feeding rate) we fit a model with a tensor product smooth interaction between current temperature and either baseline or stress induced corticosterone. In incubation models, we included data from the 7 days immediately after corticosterone measures were collected. In provisioning models, we included data from the first 7 days of incubation, which spanned the period over which corticosterone samples were collected. As in the early life analyses above, we assessed support for the interaction model by comparison to reduced models with AIC values and we plotted predictions for the 5^th^ and 95^th^ percentile of corticosterone values. We did not fit similar models for males because the sample size was much smaller due to the fact that plasma samples from males were not assayed in all years.

#### Effect of parental behavior on nestling growth and survival

To estimate the effect of parental behavioral differences on reproductive success we used the BLUPs from the initial LMMs described above as predictors of nestling survival and growth. Using BLUPs as predictors in subsequent models is known to be anti-conservative because the uncertainty in BLUP estimation is not propagated to subsequent models (*50, 51*). Therefore, in all of the models described here we created bootstrapped confidence intervals by sampling 100 BLUP estimates from the initial LMMs and then refitting the reproductive success models to generate consensus 95% confidence intervals. We interpreted results as supported only if the bootstrapped confidence interval did not cross one (odds ratio for survival models) or zero (standardized effect size for growth models).

For survival models, we fit binomial GLMMs with either hatching, survival to day 12, survival to fledge, or recruiting to the population as a breeding adult as the binomial response. In each case, predictors included the random intercept BLUP and random slope BLUP from one of the behavioral models described above (on bout duration, off bout duration, female feeding rate, or male feeding rate). Nest was included as a random effect in all models. Growth models were structured similarly to survival models, except that they were fit with a normal distribution and with average nestling mass, head size, or wing length on day 12 as predictors. Because there was only one average morphological measure per nest, no random effects were required. We scaled all response and predictor variables so that effect sizes are all interpretable in standard deviation units. We assessed the residuals and dispersion of all models using the ‘DHARMa’ package in R (*52*). In a few cases, models were overdispersed and we switched from binomial to betabinomial distribution to meet the required assumptions.

## SUPPLEMENTAL FIGURES AND TABLES

**Fig. S1.**
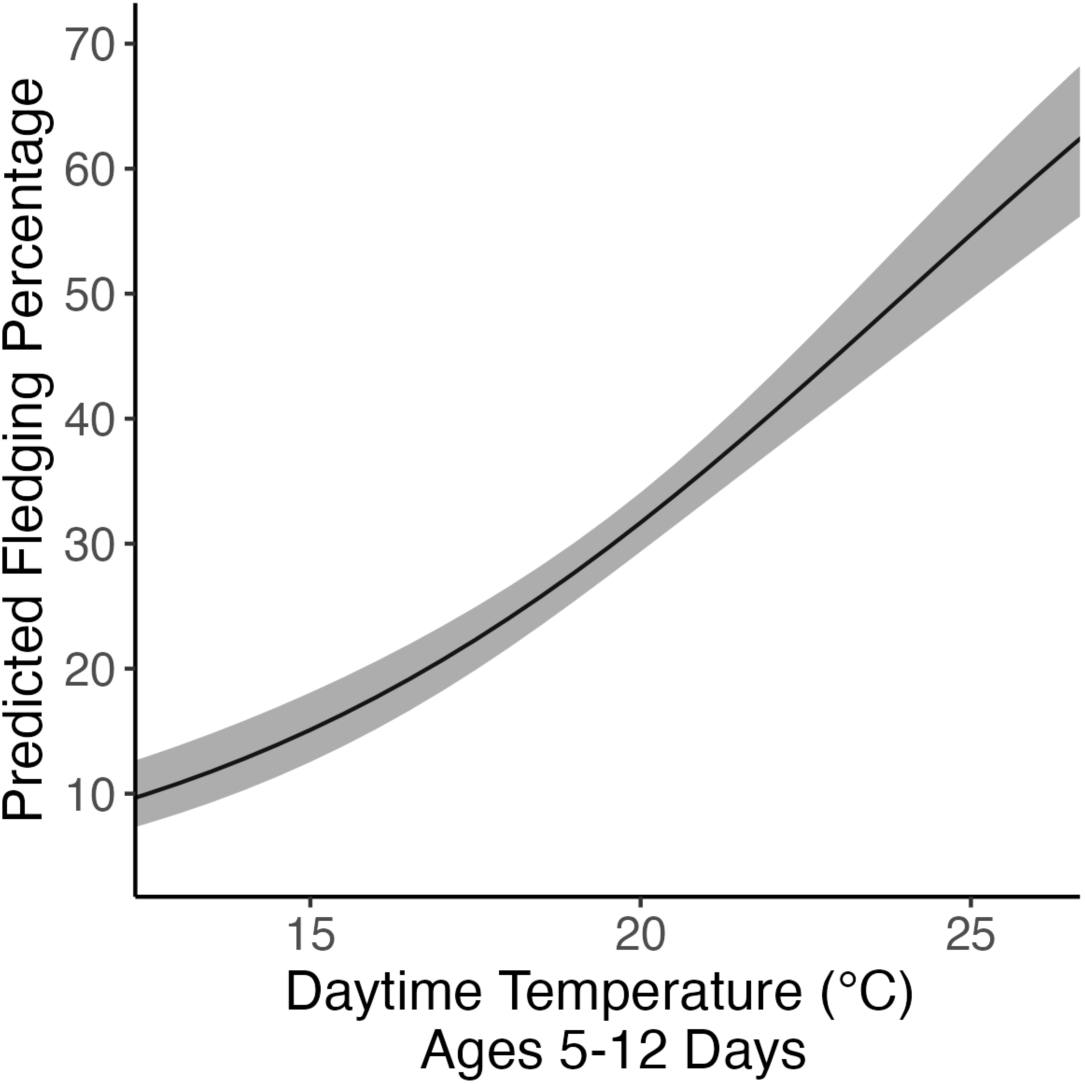
Probability of fledging in relation to temperature during the provisioning period. Figure illustrates the model predicted probability and confidence interval of fledging percentage for nestlings that hatched successfully from the long term database. Fit is from a simple binomial model with average daytime temperature for ages 5-12 days as a predictor and nest identity as a random effect (*n* = 1,837 nests and 12,353 hatched nestlings).

**Fig. S2.**
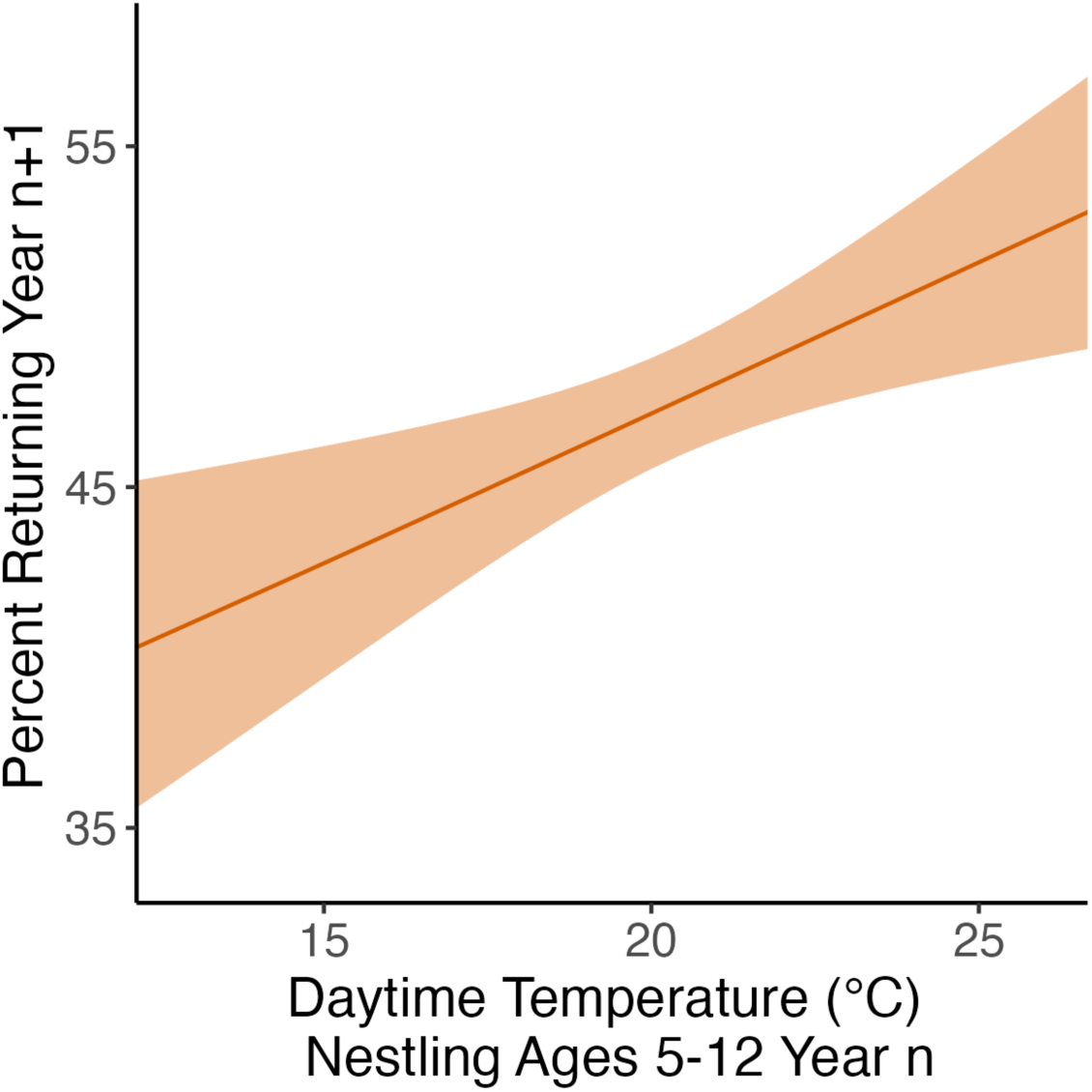
Apparent survival of adult females in relation to temperature during peak provisioning period. Figure illustrates the model predicted probability and confidence interval of returning in year n+1 for adult females from the long term database with average daytime temperature when nestlings were 5-12 days old in year n as a predictor (*n* = 3,673). Males were not included because surveys of male return were incomplete and highly variable between years.

**Fig. S3.**
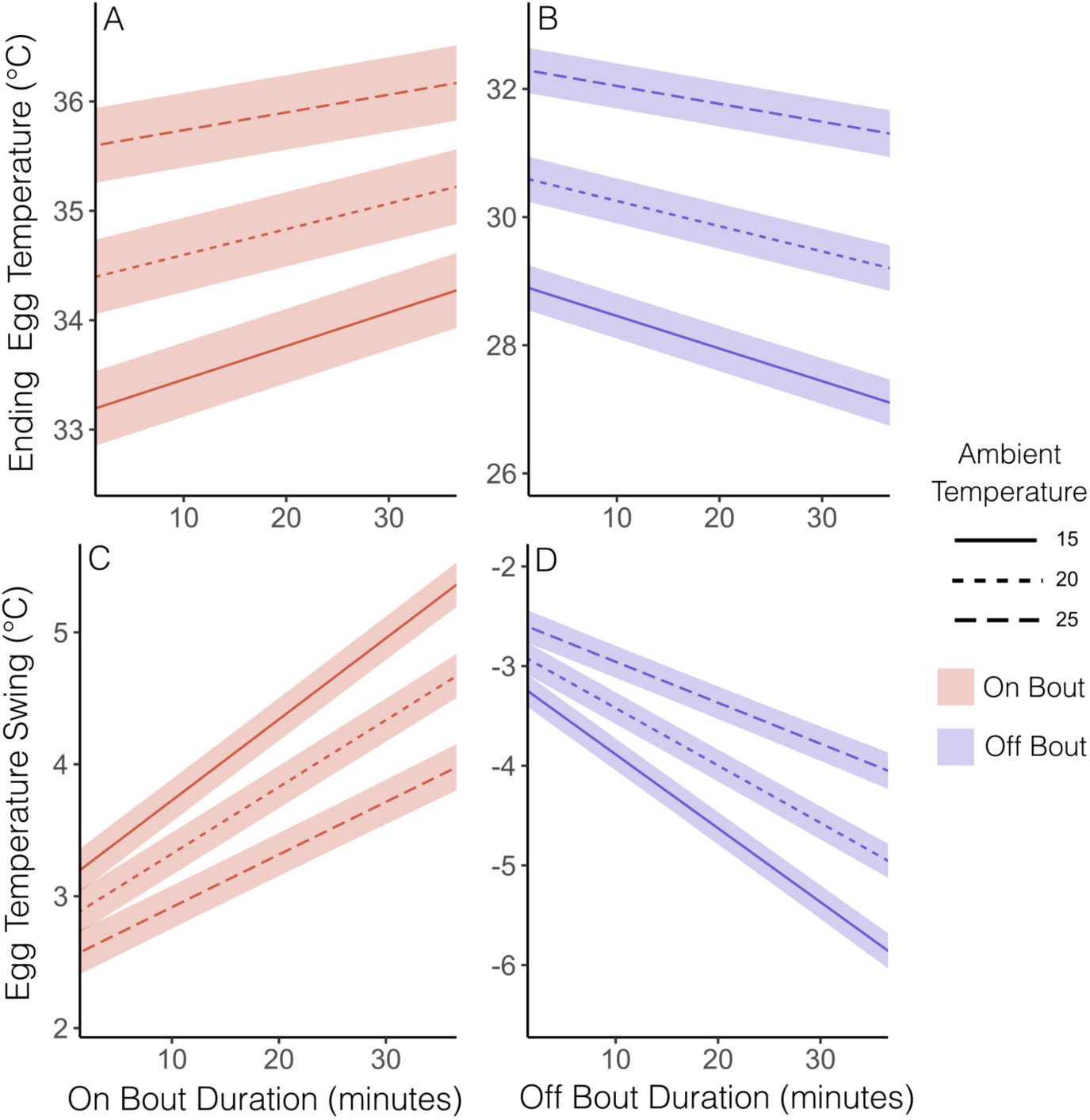
Relationship between bout duration and egg temperature at different levels of ambient temperature. Upper panels (A to B) show the temperature of the egg thermocouple at the conclusion of bouts of varying lengths at three different ambient temperature levels. Lower panels (C to D) show the change in absolute temperature of the egg thermocouple from the beginning to the end of bouts of varying length at three different ambient temperature levels.

**Fig. S4.**
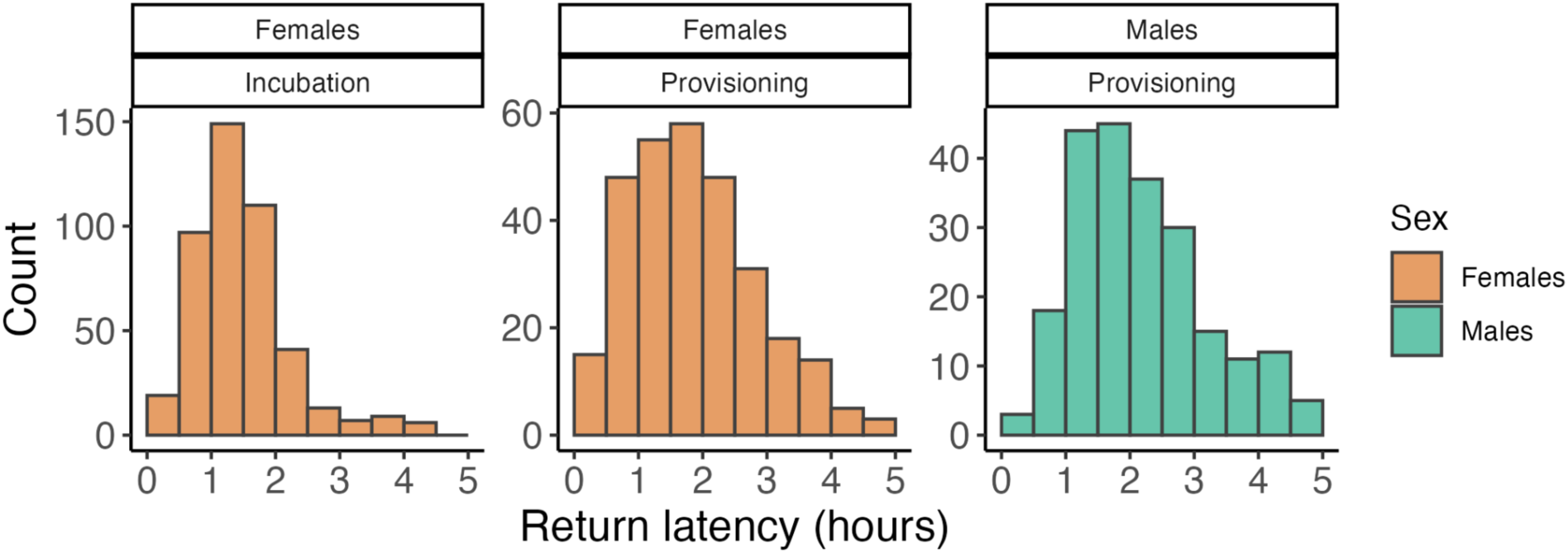
Variation in latency to return to the nest box after capture. Latency to return was determined by the elapsed time from release to the first RFID reading of the focal bird at their home nest box. Panels illustrate the distribution of return latency times for females during incubation or provisioning and for males during provisioning.

**Fig. S5.**
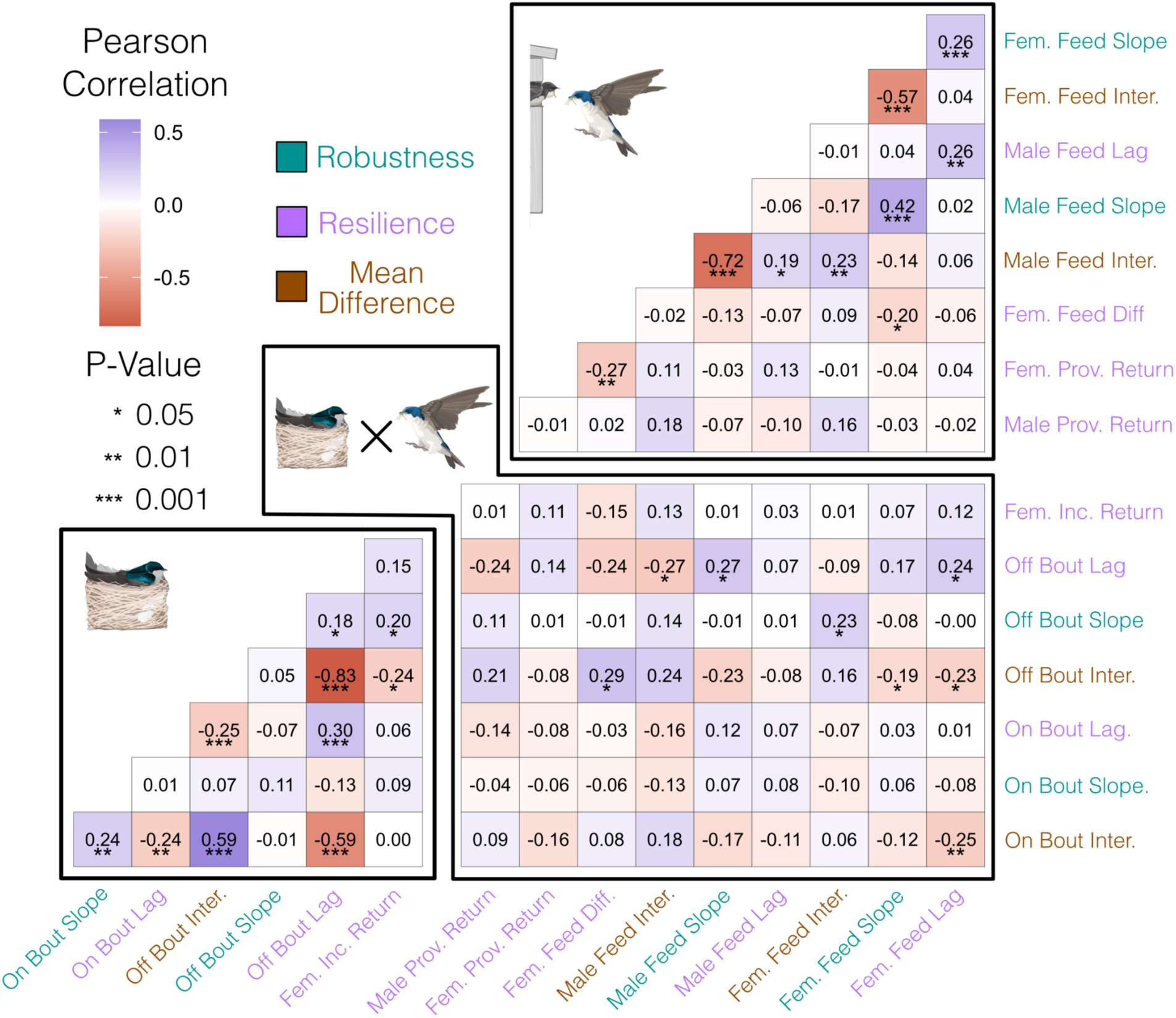
Relationship between each behavioral measure in the study. Pearson correlations for each pair of measures. Lower triangle denotes behavioral measures extracted from incubation data. Upper triangle denotes behavioral measures extracted from provisioning data. Rectangle illustrates relationships between pairs of incubation vs. provisioning behavioral data. Sample size differed for each comparison based on available data. Illustrations by Charlotte Holden, reproduced with permission from the Cornell Lab of Ornithology.

**Fig. S6.**
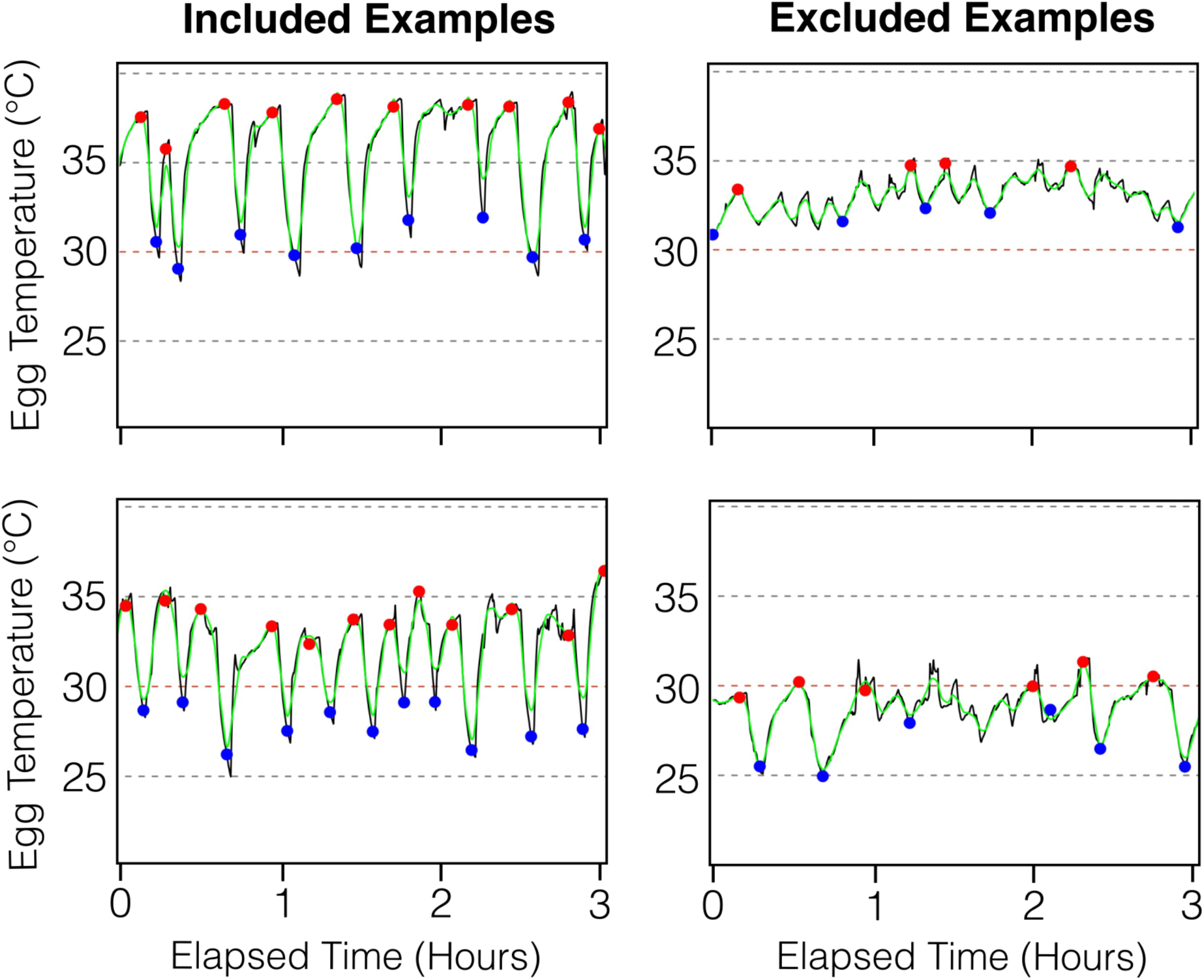
Illustration of thermocouple processing script and manual filtering to exclude sections that could not be scored. In each panel black line shows the raw data values, green line shows the smoothed curve fit to the data, and dots show the ending point of off bouts (blue) and on bouts (red) identified by the algorithm. The left panels illustrate examples of good sections that would have passed visual inspection for inclusion. Note that in the lower left panel, the thermocouple may not be positioned well to record precise absolute temperature, but the timing of on and off bouts is still clear. The right panels illustrate two sections that would have been excluded from scoring. Despite some cyclical temperature changes, there is no clear identification of on and off bouts, perhaps due to poor thermocouple placement or reading errors. In many cases, excluded sections would include a flat line mirroring ambient temperature if the thermocouple was entirely out of the nest. All excluded sections were filtered out of the final data before analyses.

**Table S1.** Effect of temperature on adult mass and nestling morphology. Details of generalized additive mixed models including one measure as a response with day relative to hatching and temperature as smoothed predictors. Models were fit with gamm4 and also include a random effect for individual identity. For each response variable, models are ranked by AIC score. Partial effects from models correspond to plots in Fig. 1.

| *Predictor* | *edf* | *F* | *P* | $\Delta AIC$ |
| --- | --- | --- | --- | --- |
| **Adult Female Mass** |  |  |  |  |
| 3-Day Temperature ($n = 7,843$ measures 3,669 individuals; Cond. $R^2 = 0.58$; Adj $R^2 = 0.42$) | | | | |
| Days From Hatch | 7.83 | 889.60 | < 0.001 | 0 |
| 3-Day Temperature | 7.68 | 313.10 | < 0.001 |  |
| 5-Day Temperature ($n = 7,843$ measures 3,669 individuals; Cond. $R^2 = 0.57$; Adj $R^2 = 0.39$) | | | | |
| Days From Hatch | 7.82 | 803.20 | < 0.001 | 492.06 |
| 5-Day Temperature | 7.19 | 242.90 | < 0.001 |  |
| 1-Day Temperature ($n = 7,843$ measures 3,669 individuals; Cond. $R^2 = 0.45$; Adj $R^2 = 0.38$) | | | | |
| Days From Hatch | 7.93 | 769.20 | < 0.001 | 713.12 |
| 1-Day Temperature | 6.19 | 233.40 | < 0.001 |  |
| **Adult Male Mass** |  |  |  |  |
| 1-Day Temperature ($n = 2,731$ measures 1,898 individuals; Cond. $R^2 = 0.51$; Adj $R^2 = 0.05$) | | | | |
| Days From Hatch | 4.05 | 40.67 | < 0.001 | 0 |
| 1-Day Temperature | 2.10 | 39.98 | < 0.001 |  |
| 3-Day Temperature ($n = 2,731$ measures 1,898 individuals; Cond. $R^2 = 0.51$; Adj $R^2 = 0.05$) | | | | |
| Days From Hatch | 3.76 | 45.31 | < 0.001 | 16.68 |
| 3-Day Temperature | 2.52 | 26.32 | < 0.001 |  |
| 5-Day Temperature ($n = 2,731$ measures 1,898 individuals; Cond. $R^2 = 0.51$; Adj $R^2 = 0.05$) | | | | |
| Days From Hatch | 4.10 | 38.38 | < 0.001 | 59.25 |
| 5-Day Temperature | 1.26 | 17.41 | < 0.001 |  |
| **Nestling Mass** |  |  |  |  |
| 3-Day Temperature ($n = 21,049$ measures 3,506 nests; Cond. $R^2 = 0.68$; Adj $R^2 = 0.67$) | | | | |
| Days From Hatch | 8.55 | 718.42 | < 0.001 | 0 |
| 3-Day Temperature | 13.63 | 25.21 | < 0.001 |  |
| 5-Day Temperature ($n = 21,049$ measures 3,506 nests; Cond. $R^2 = 0.71$; Adj $R^2 = 0.67$) | | | | |
| Days From Hatch | 8.50 | 308.88 | < 0.001 | 37.4 |
| 5-Day Temperature | 15.32 | 19.39 | < 0.001 |  |
| 1-Day Temperature ($n = 21,049$ measures 3,506 nests; Cond. $R^2 = 0.68$; Adj $R^2 = 0.66$) | | | | |
| Days From Hatch | 8.59 | 648.80 | < 0.001 | 303.43 |
| 1-Day Temperature | 16.09 | 16.09 | < 0.001 |  |
| **Nestling Flattened Wing Length** |  |  |  |  |
| 5-Day Temperature ($n = 18,469$ measures 3,315 nests; Cond. $R^2 = 0.77$; Adj $R^2 = 0.72$) | | | | |
| Days From Hatch | 7.14 | 299.06 | < 0.001 | 0 |
| 5-Day Temperature | 15.94 | 24.67 | < 0.001 |  |
| 3-Day Temperature ($n = 18,469$ measures 3,315 nests; Cond. $R^2 = 0.80$; Adj $R^2 = 0.70$) | | | | |
| Days From Hatch | 7.72 | 338.52 | < 0.001 | 269.4 |
| 3-Day Temperature | 17.12 | 37.79 | < 0.001 |  |
| 1-Day Temperature ($n = 18,469$ measures 3,315 nests; Cond. $R^2 = 0.84$; Adj $R^2 = 0.69$) | | | | |
| Days From Hatch | 8.13 | 332.80 | < 0.001 | 649.5 |
| 1-Day Temperature | 16.90 | 15.60 | < 0.001 |  |
| **Nestling Head + Bill Length** |  |  |  |  |
| 5-Day Temperature ($n = 15,336$ measures 2,650 nests; Cond. $R^2 = 0.81$; Adj $R^2 = 0.82$) | | | | |
| Days From Hatch | 7.81 | 611.40 | < 0.001 | 0 |
| 5-Day Temperature | 12.41 | 9.67 | < 0.001 |  |
| 3-Day Temperature ($n = 15,336$ measures 2,650 nests; Cond. $R^2 = 0.83$; Adj $R^2 = 0.82$) | | | | |
| Days From Hatch | 7.58 | 1327.40 | < 0.001 | 70.55 |
| 3-Day Temperature | 12.18 | 11.73 | < 0.001 |  |
| 1-Day Temperature ($n = 15,336$ measures 2,650 nests; Cond. $R^2 = 0.82$; Adj $R^2 = 0.82$) | | | | |
| Days From Hatch | 7.83 | 1436.92 | < 0.001 | 385.78 |
| 1-Day Temperature | 10.80 | 12.07 | < 0.001 |  |

**Table S2.** Description of all behavioral traits extracted from automated monitoring systems. Each trait in the table was measured for each individual with appropriate data, thus sample sizes and overlap in measures available varies by individual.

| Stage | Behavior | Type | Description, Interpretation, and Measurement |
| --- | --- | --- | --- |
| Inc. | On/Off Bout Length | Individual Difference | The typical length of on or off bouts accounting for ambient temperature. Measured as the random effect intercept BLUP from the on or off bout model. Females only. |
| Inc. | On/Off Bout Sensitivity | Robustness | The sensitivity of on or off bout length to declining temperature. Measured as the random effect slope BLUP from the on bout model. A steeper negative slope (lower value) is interpreted as lower robustness to temperature. Females only. |
| Inc. | On/Off Bout Recovery | Resilience | Recovery of incubation bouts to normal length with respect to prior temperature. Measured as the random slope BLUP from the lagged temperature model. |
| Inc. & Prov. | Latency to Return | Resilience | Latency to return to the nest box after capture during incubation or provisioning. Measured from RFID records as the time between release and first reading at the focal nest box. Faster return is interpreted as higher resilience. Females at both stages; males only in provisioning. |
| Prov. | Feeding Rate | Individual Difference | The individual hourly feeding rate accounting for ambient temperature. Measured as the random effect intercept BLUP from the feeding model. Both sexes. |
| Prov. | Feeding Sensitivity | Robustness | The sensitivity of hourly feeding rate to declining temperature. Measured as the random effect slope BLUP from the feeding model. Steeper positive slope (higher value) interpreted as lower robustness. Both sexes. |
| Prov. | Feeding Difference | Resilience | Change in feeding rate following a major disturbance. Measured as change in average feeding rate on the afternoon before capture vs. the afternoon on the day of capture (with capture in early morning). Negative values interpreted as lower resilience. Females only. |
| Prov. | Feeding Recovery | Resilience | Recovery of feeding to normal rates with respect to prior temperature. Measured as the random slope BLUP from a lagged temperature model. Steeper positive slope interpreted as lower resilience. Both sexes. |

**Table S3.** Relationship between on or off bout length and absolute temperature of eggs during incubation. All variables are standardized to mean of zero and standard deviation of one. All models include nest identity as a random effect.

| *Parameter* | *Estimate* | *CI* | *t-value* | *P-value* |
| --- | --- | --- | --- | --- |
| **On Bout Duration ($n = 44,220$ bouts from 209 nests)** |  |  |  |  |
| **Maximum egg temperature (marginal $R^2 = 0.12$; conditional $R^2 = 0.77$)** |  |  |  |  |
| Intercept | -0.09 | [-0.21, 0.02] | -1.60 | 0.11 |
| Bout Duration | 0.08 | [0.07, 0.08] | 29.8 | <0.0001 |
| Ambient Temperature | 0.37 | [0.36, 0.37] | 134.6 | <0.0001 |
| Duration * Temperature | -0.02 | [-0.03, -0.02] | -10.43 | <0.0001 |
| **Temperature swing during bout (marginal $R^2 = 0.09$; conditional $R^2 = 0.34$)** |  |  |  |  |
| Intercept | -0.11 | [-0.19, -0.04] | -3.20 | 0.002 |
| Bout Duration | 0.21 | [0.21, 0.22] | 50.8 | <0.0001 |
| Ambient Temperature | -0.19 | [-0.20, -0.18] | -42.1 | <0.0001 |
| Duration * Temperature | -0.04 | [-0.05, -0.04] | -12.4 | <0.0001 |
| **Off Bout Duration ($n = 44834$ bouts from 209 nests)** |  |  |  |  |
| **Minimum egg temperature (marginal $R^2 = 0.23$; conditional $R^2 = 0.71$)** |  |  |  |  |
| Intercept | 0.01 | [-0.09, 0.11] | 0.20 | 0.83 |
| Bout Duration | -0.09 | [-0.09, -0.08] | -30.71 | <0.0001 |
| Ambient Temperature | 0.50 | [0.49, 0.50] | 163.7 | <0.0001 |
| Duration * Temperature | 0.02 | [0.02, 0.03] | 10.2 | <0.0001 |
| **Temperature swing during bout (marginal $R^2 = 0.09$; conditional $R^2 = 0.34$)** |  |  |  |  |
| Intercept | 0.11 | [0.04, 0.18] | 0.30 | 0.002 |
| Bout Duration | -0.20 | [-0.21, -0.19] | -48.8 | <0.0001 |
| Ambient Temperature | 0.21 | [0.20, 0.21] | 46.1 | <0.0001 |
| Duration * Temperature | 0.06 | [0.05, 0.06] | 16 | <0.0001 |

**Table S4.**
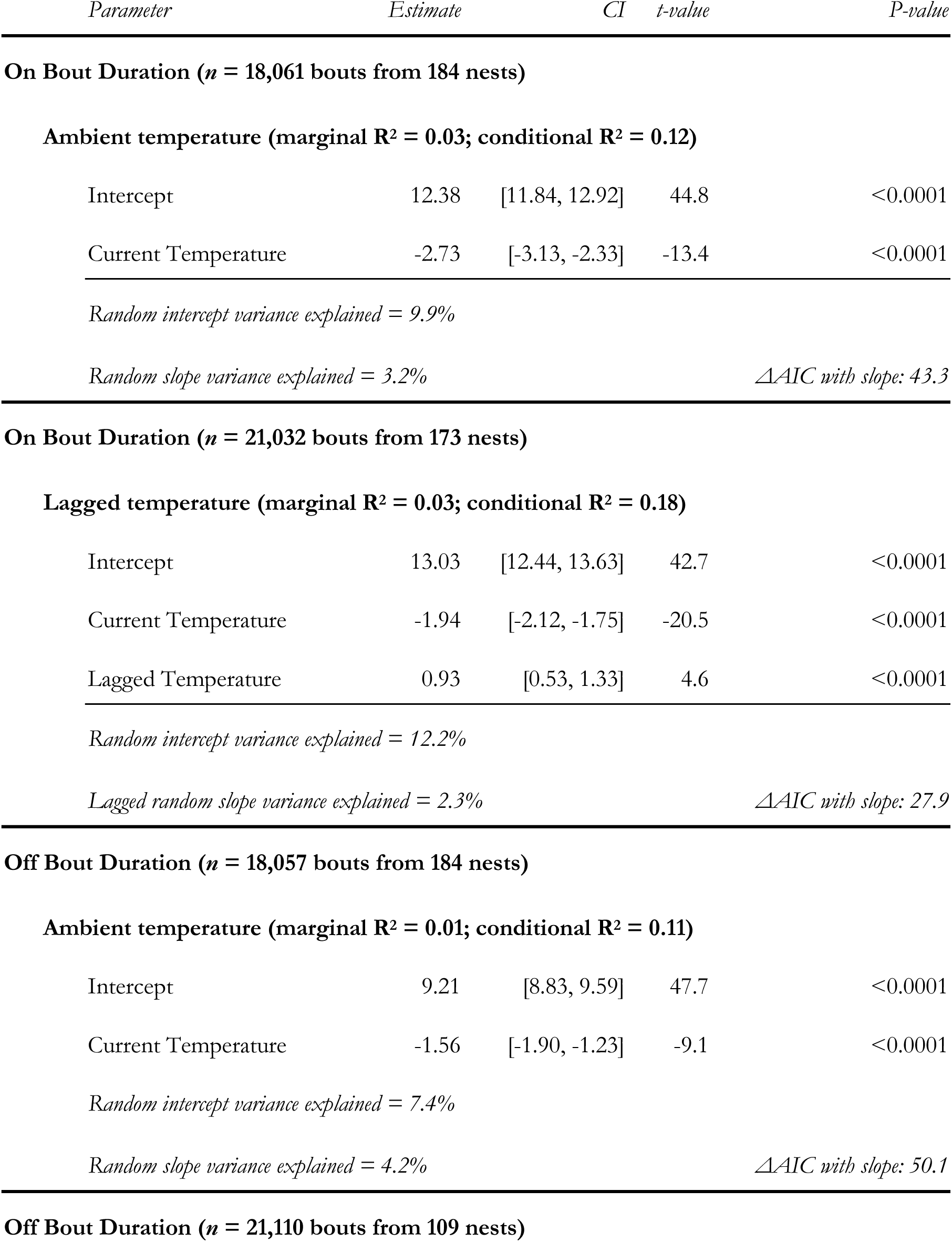
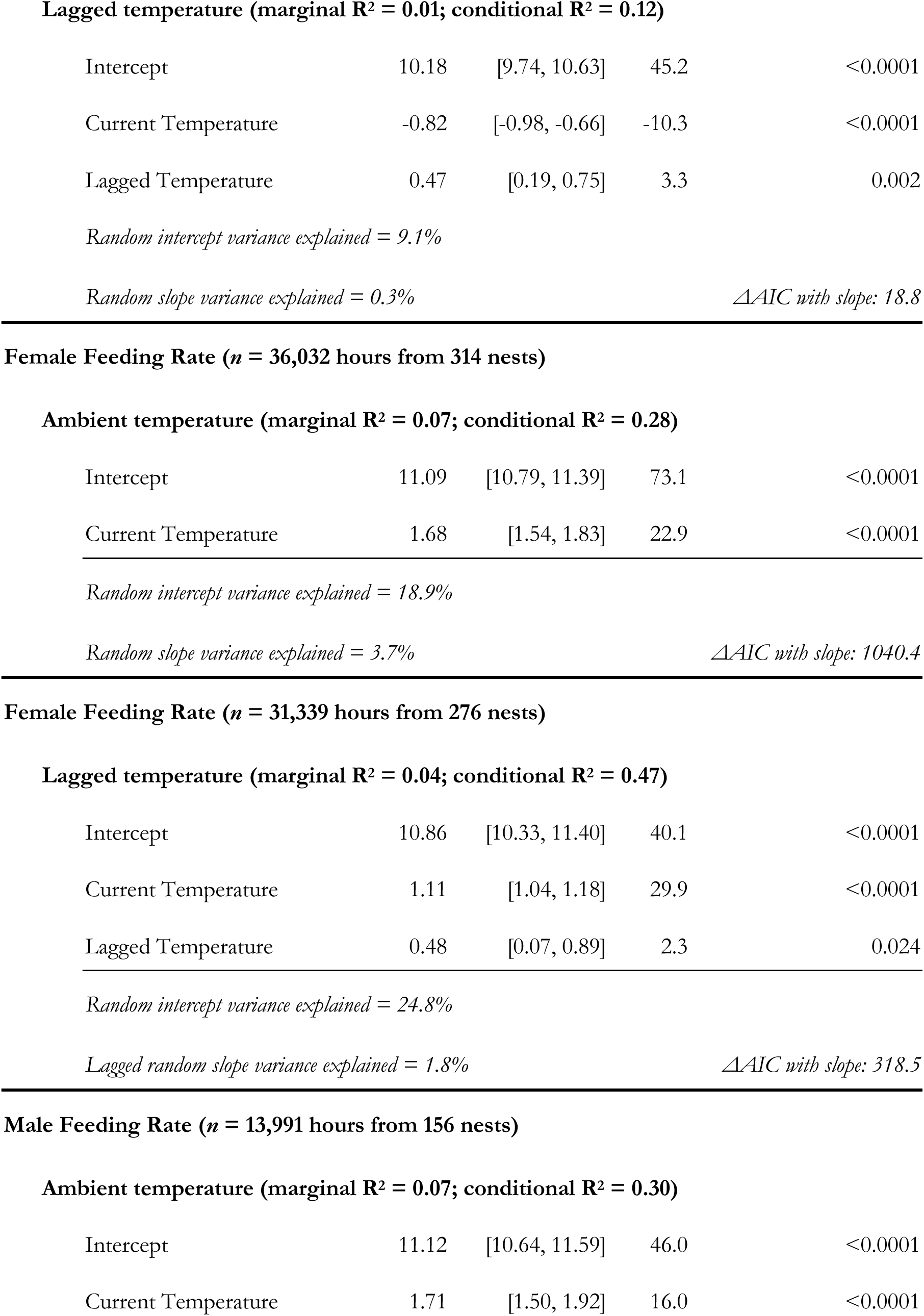
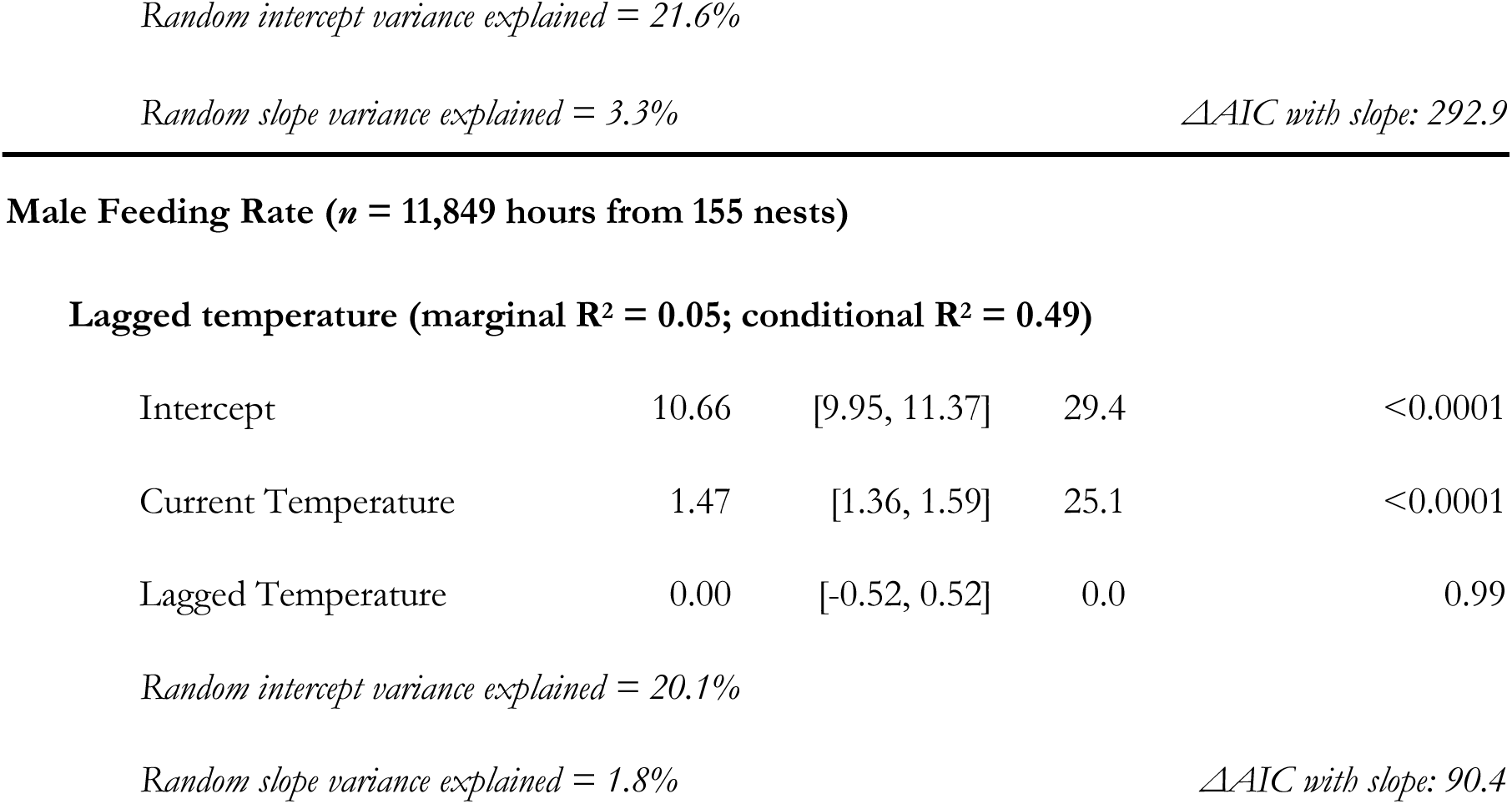
Models of individual differences in behavioral sensitivity to declining temperature. For each response variable (on or off bout length and male or female feeding rate), we fit one model with a random slope to current hourly temperature and a separate model with a random slope to the prior day’s temperature accounting for current temperature.

| *Parameter* | *Estimate* | *CI* | *t-value* | *P-value* |
| --- | --- | --- | --- | --- |
| **On Bout Duration (*n* = 18,061 bouts from 184 nests)** |  |  |  |  |
| **Ambient temperature (marginal R<sup>2</sup> = 0.03; conditional R<sup>2</sup> = 0.12)** |  |  |  |  |
| Intercept | 12.38 | [11.84, 12.92] | 44.8 | <0.0001 |
| Current Temperature | -2.73 | [-3.13, -2.33] | -13.4 | <0.0001 |
| *Random intercept variance explained* = 9.9% |  |  |  |  |
| *Random slope variance explained* = 3.2% |  |  |  | *ΔAIC with slope: 43.3* |
| **On Bout Duration (*n* = 21,032 bouts from 173 nests)** |  |  |  |  |
| **Lagged temperature (marginal R<sup>2</sup> = 0.03; conditional R<sup>2</sup> = 0.18)** |  |  |  |  |
| Intercept | 13.03 | [12.44, 13.63] | 42.7 | <0.0001 |
| Current Temperature | -1.94 | [-2.12, -1.75] | -20.5 | <0.0001 |
| Lagged Temperature | 0.93 | [0.53, 1.33] | 4.6 | <0.0001 |
| *Random intercept variance explained* = 12.2% |  |  |  |  |
| *Lagged random slope variance explained* = 2.3% |  |  |  | *ΔAIC with slope: 27.9* |
| **Off Bout Duration (*n* = 18,057 bouts from 184 nests)** |  |  |  |  |
| **Ambient temperature (marginal R<sup>2</sup> = 0.01; conditional R<sup>2</sup> = 0.11)** |  |  |  |  |
| Intercept | 9.21 | [8.83, 9.59] | 47.7 | <0.0001 |
| Current Temperature | -1.56 | [-1.90, -1.23] | -9.1 | <0.0001 |
| *Random intercept variance explained* = 7.4% |  |  |  |  |
| *Random slope variance explained* = 4.2% |  |  |  | *ΔAIC with slope: 50.1* |
| **Off Bout Duration (*n* = 21,110 bouts from 109 nests)** |  |  |  |  |
| **Lagged temperature (marginal $R^2 = 0.01$ ; conditional $R^2 = 0.12$ )** |  |  |  |  |
| Intercept | 10.18 | [9.74, 10.63] | 45.2 | <0.0001 |
| Current Temperature | -0.82 | [-0.98, -0.66] | -10.3 | <0.0001 |
| Lagged Temperature | 0.47 | [0.19, 0.75] | 3.3 | 0.002 |
| Intercept | 11.09 | [10.79, 11.39] | 73.1 | <0.0001 |
| Current Temperature | 1.68 | [1.54, 1.83] | 22.9 | <0.0001 |
| Intercept | 10.86 | [10.33, 11.40] | 40.1 | <0.0001 |
| Current Temperature | 1.11 | [1.04, 1.18] | 29.9 | <0.0001 |
| Lagged Temperature | 0.48 | [0.07, 0.89] | 2.3 | 0.024 |
| Intercept | 11.12 | [10.64, 11.59] | 46.0 | <0.0001 |
| Current Temperature | 1.71 | [1.50, 1.92] | 16.0 | <0.0001 |
| **Lagged temperature (marginal $R^2 = 0.05$ ; conditional $R^2 = 0.49$ )** |  |  |  |  |
| Intercept | 10.66 | [9.95, 11.37] | 29.4 | <0.0001 |
| Current Temperature | 1.47 | [1.36, 1.59] | 25.1 | <0.0001 |
| Lagged Temperature | 0.00 | [-0.52, 0.52] | 0.0 | 0.99 |

*Random intercept variance explained = 20.1%*
*Random slope variance explained = 1.8%*
*ΔAIC with slope: 90.4*

**Table S5.** Effect size estimates for behavioral measures as predictors of nestling hatching, survival to day 12, survival to fledging, and recruitment in following year. Estimates are reported as odds ratios from binomial models. 95% confidence intervals are reported from the initial model and from 100 bootstrapped models fit to account for uncertainty in BLUP estimation. Confidence intervals that do not cross 1 are shown in bold.

| Response | Predictor | n chicks | n nests | Effect Size | Conf. Interval | Boot Interval |
| --- | --- | --- | --- | --- | --- | --- |
| Hatch | On Bout Intercept | 943 | 170 | 1.92 | **[1.23, 2.97]** | **[1.01, 3.27]** |
|  | On Bout Slope | 943 | 170 | 0.69 | [0.45, 1.06] | [0.41, 1.62] |
|  | Off Bout Intercept | 943 | 170 | 1.64 | **[1.05, 2.56]** | [0.68, 3.40] |
|  | Off Bout Slope | 943 | 170 | 1.00 | [0.65, 1.54] | [0.44, 2.25] |
| Day 12 | On Bout Intercept | 777 | 170 | 1.85 | **[1.33, 2.57]** | **[1.07, 2.52]** |
|  | On Bout Slope | 777 | 170 | 0.79 | [0.59, 1.06] | [0.51, 1.34] |
|  | Off Bout Intercept | 777 | 170 | 1.28 | [0.94, 1.74] | [0.74, 2.46] |
|  | Off Bout Slope | 777 | 170 | 1.16 | [0.88, 1.54] | [0.64, 2.22] |
|  | Fem Feed Slope | 1516 | 314 | 2.36 | **[1.74, 3.19]** | **[1.48, 3.38]** |
|  | Fem Feed Intercept | 1516 | 314 | 0.65 | [0.49, 0.85] | [0.44, 1.01] |
|  | Male Feed Slope | 762 | 156 | 3.01 | **[1.75, 5.16]** | **[1.16, 6.73]** |
|  | Male Feed Intercept | 762 | 156 | 0.67 | [0.41, 1.10] | [0.31, 1.67] |
| Fledge | On Bout Intercept | 774 | 169 | 1.40 | [0.83, 2.36] | [0.61, 2.93] |
|  | On Bout Slope | 774 | 169 | 0.77 | [0.46, 1.31] | [0.30, 2.07] |
|  | Off Bout Intercept | 774 | 169 | 1.21 | [0.74, 1.98] | [0.56, 3.64] |
|  | Off Bout Slope | 774 | 169 | 1.57 | [0.97, 2.55] | [0.59, 4.99] |
|  | Fem Feed Slope | 1512 | 313 | 2.32 | **[1.80, 3.00]** | **[1.54, 3.26]** |
|  | Fem Feed Intercept | 1512 | 313 | 0.93 | [0.74, 1.16] | [0.65, 1.31] |
|  | Male Feed Slope | 758 | 155 | 1.96 | **[1.31, 2.95]** | [0.96, 3.59] |
|  | Male Feed Intercept | 758 | 155 | 0.71 | [0.49, 1.03] | [0.38, 1.46] |
| Recruit | On Bout Intercept | 442 | 169 | 1.01 | [0.56, 1.83] | [0.47, 2.24] |
|  | On Bout Slope | 442 | 169 | 0.82 | [0.45, 1.49] | [0.27, 2.04] |
|  | Off Bout Intercept | 442 | 169 | 1.06 | [0.59, 1.89] | [0.37, 3.69] |
|  | Off Bout Slope | 442 | 169 | 1.35 | [0.67, 2.70] | [0.32, 4.53] |
|  | Fem Feed Slope | 888 | 313 | 2.14 | **[1.16, 3.93]** | [0.93, 4.72] |
|  | Fem Feed Intercept | 888 | 313 | 0.70 | [0.43, 1.15] | [0.34, 1.49] |
|  | Male Feed Slope | 481 | 155 | 1.51 | [0.81, 2.81] | [0.61, 4.61] |
|  | Male Feed Intercept | 481 | 155 | 0.60 | [0.32, 1.11] | [0.19, 1.72] |

**Table S6.**
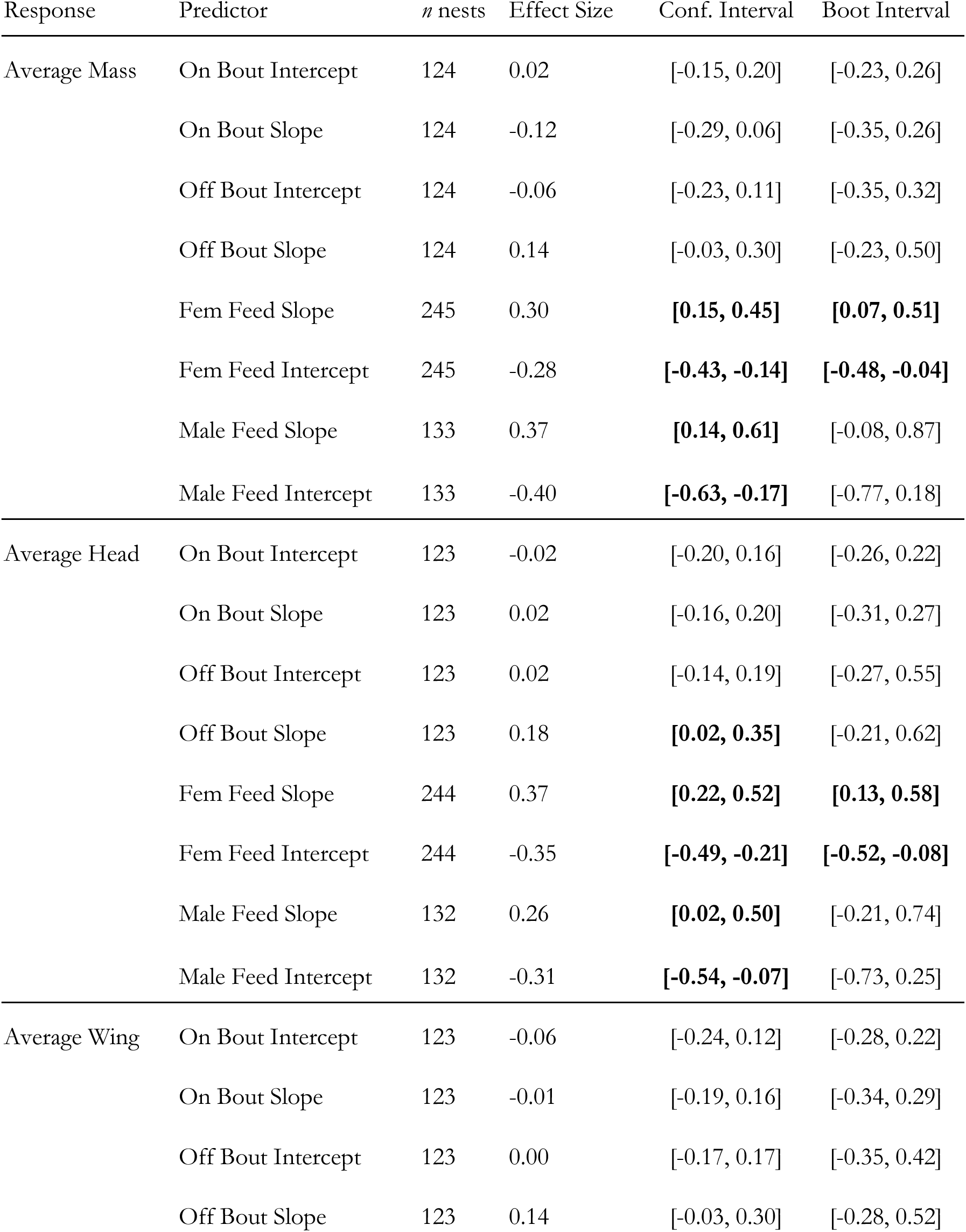
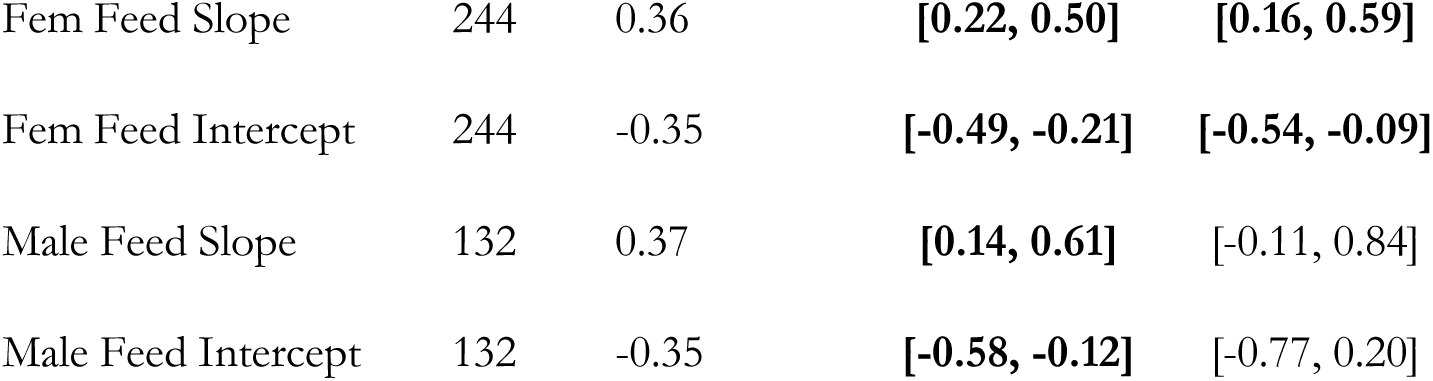
Effect size estimates for behavioral measures as predictors of average nestling mass, head plus bill length, and flattened wing length at 12 days old. Estimates are reported as standardized effect sizes. 95% confidence intervals are reported from the initial model and from 100 bootstrapped models fit to account for uncertainty in BLUP estimation. Confidence intervals that do not cross 0 are shown in bold.

| Response | Predictor | *n* nests | Effect Size | Conf. Interval | Boot Interval |
| --- | --- | --- | --- | --- | --- |
| Average Mass | On Bout Intercept | 124 | 0.02 | [-0.15, 0.20] | [-0.23, 0.26] |
|  | On Bout Slope | 124 | -0.12 | [-0.29, 0.06] | [-0.35, 0.26] |
|  | Off Bout Intercept | 124 | -0.06 | [-0.23, 0.11] | [-0.35, 0.32] |
|  | Off Bout Slope | 124 | 0.14 | [-0.03, 0.30] | [-0.23, 0.50] |
|  | Fem Feed Slope | 245 | 0.30 | **[0.15, 0.45]** | **[0.07, 0.51]** |
|  | Fem Feed Intercept | 245 | -0.28 | **[-0.43, -0.14]** | **[-0.48, -0.04]** |
|  | Male Feed Slope | 133 | 0.37 | **[0.14, 0.61]** | [-0.08, 0.87] |
|  | Male Feed Intercept | 133 | -0.40 | **[-0.63, -0.17]** | [-0.77, 0.18] |
| Average Head | On Bout Intercept | 123 | -0.02 | [-0.20, 0.16] | [-0.26, 0.22] |
|  | On Bout Slope | 123 | 0.02 | [-0.16, 0.20] | [-0.31, 0.27] |
|  | Off Bout Intercept | 123 | 0.02 | [-0.14, 0.19] | [-0.27, 0.55] |
|  | Off Bout Slope | 123 | 0.18 | **[0.02, 0.35]** | [-0.21, 0.62] |
|  | Fem Feed Slope | 244 | 0.37 | **[0.22, 0.52]** | **[0.13, 0.58]** |
|  | Fem Feed Intercept | 244 | -0.35 | **[-0.49, -0.21]** | **[-0.52, -0.08]** |
|  | Male Feed Slope | 132 | 0.26 | **[0.02, 0.50]** | [-0.21, 0.74] |
|  | Male Feed Intercept | 132 | -0.31 | **[-0.54, -0.07]** | [-0.73, 0.25] |
| Average Wing | On Bout Intercept | 123 | -0.06 | [-0.24, 0.12] | [-0.28, 0.22] |
|  | On Bout Slope | 123 | -0.01 | [-0.19, 0.16] | [-0.34, 0.29] |
|  | Off Bout Intercept | 123 | 0.00 | [-0.17, 0.17] | [-0.35, 0.42] |
|  | Off Bout Slope | 123 | 0.14 | [-0.03, 0.30] | [-0.28, 0.52] |

|  |  |  |  |  |
| --- | --- | --- | --- | --- |
| Fem Feed Slope | 244 | 0.36 | **[0.22, 0.50]** | **[0.16, 0.59]** |
| Fem Feed Intercept | 244 | -0.35 | **[-0.49, -0.21]** | **[-0.54, -0.09]** |
| Male Feed Slope | 132 | 0.37 | **[0.14, 0.61]** | [-0.11, 0.84] |
| Male Feed Intercept | 132 | -0.35 | **[-0.58, -0.12]** | [-0.77, 0.20] |

**Table S7.** Comparison of models exploring the effect of early life incubation temperature or current corticosterone expression on incubation bout duration and provisioning rate. Models were fit as generalized linear mixed models with smoothed interactions and compared to simplified models that included either separate smooths for the two predictors or only current temperature. All models included a random effect for nest identity.

| *Model* | *Predictors* | *k* | *weight* | *ΔAIC* |
| --- | --- | --- | --- | --- |
| **Early Life Conditions** |  |  |  |  |
| Feeding Rate (n = 7271 hours from 54 females) |  |  |  |  |
| 1 | Temperature * Natal Incubation Temperature | 9 | 1.00 | 0.00 |
| 3 | Temperature | 5 | <0.001 | 50.15 |
| 2 | Temperature + Natal Incubation Temperature | 7 | <0.001 | 57.64 |
| **Current Corticosterone State** |  |  |  |  |
| Incubation On Bout Duration & Baseline Corticosterone (9,069 bouts from 71 nests) |  |  |  |  |
| 3 | Temperature | 5 | 0.64 | 0.00 |
| 2 | Temperature + Baseline Corticosterone | 7 | 0.24 | 1.97 |
| 1 | Temperature * Baseline Corticosterone | 9 | 0.13 | 3.59 |
| Incubation On Bout Duration & Stress-Induced Corticosterone (9,069 bouts from 71 nests) |  |  |  |  |
| 3 | Temperature | 5 | 0.68 | 0.00 |
| 2 | Temperature + Stress-Induced Corticosterone | 7 | 0.26 | 2.53 |
| 1 | Temperature * Stress-Induced Corticosterone | 9 | 0.06 | 9.15 |
| Incubation Off Bout Duration & Baseline Corticosterone (9,153 bouts from 71 nests) |  |  |  |  |
| 3 | Temperature | 5 | 0.80 | 0.00 |
| 2 | Temperature + Baseline Corticosterone | 7 | 0.11 | 3.92 |
| 1 | Temperature * Baseline Corticosterone | 9 | 0.09 | 8.93 |
| Incubation Off Bout Duration & Stress-Induced Corticosterone (9,153 bouts from 71 nests) |  |  |  |  |
| 3 | Temperature | 5 | 0.80 | 0.00 |
| 2 | Temperature + Stress-Induced Corticosterone | 7 | 0.13 | 3.71 |
| 1 | Temperature * Stress-Induced Corticosterone | 9 | 0.07 | 6.30 |
| Provisioning Rate & Baseline Corticosterone (8,399 hours from 147 nests) |  |  |  |  |
| 1 | Temperature * Baseline Corticosterone | 9 | 0.84 | 0.00 |
| 3 | Temperature | 5 | 0.12 | 25.56 |
| 2 | Temperature + Baseline Corticosterone | 7 | 0.05 | 27.18 |
| Provisioning Rate & Stress-Induced Corticosterone (7,480 hours from 132 nests) |  |  |  |  |
| 3 | Temperature | 5 | 0.64 | 0.00 |
| 2 | Temperature + Baseline Corticosterone | 7 | 0.29 | 2.53 |
| 1 | Temperature * Baseline Corticosterone | 9 | 0.08 | 13.39 |

